# Comprehensive analysis of pseudogene expression in human and macaque brains compared with other tissues

**DOI:** 10.1101/2025.09.02.673826

**Authors:** Yunzhe Jiang, Yucheng T. Yang, Cristina Sisu, Tianxing He, Hyejung Won, Mark Gerstein

**Affiliations:** Program in Computational Biology and Biomedical Informatics, Yale University, New Haven, CT 06520, USA; Department of Molecular Biophysics and Biochemistry, Yale University, New Haven, CT 06520, USA; Department of Life Sciences, Brunel University of London, Uxbridge, London, UB8 3PH, UK; Department of Electrical Engineering, Columbia University, New York, NY 10027, USA; UNC Neuroscience Center, University of North Carolina, Chapel Hill, NC 27599, USA; Department of Genetics, University of North Carolina, Chapel Hill, NC 27599, USA; Department of Computer Science, Yale University, New Haven, CT 06520, USA; Department of Statistics and Data Science, Yale University, New Haven, CT 06520, USA; Department of Biomedical Informatics & Data Science, Yale University, New Haven, CT 06520, USA

## Abstract

Although gene expression in the brain has been extensively investigated and compared to other tissues, the activity of pseudogenes has not been comprehensively surveyed. Here, leveraging large-scale RNA-seq data, we construct consistent pseudogene expression profiles in human and macaque brains and compare them to 29 other tissues. We further annotate pseudogenes with potential cellular roles based on co-clustering them with protein-coding genes. Notably, the majority of the expressed pseudogenes show elevated expression in the brain relative to other tissues, and these pseudogenes show broad and consistent expression patterns across brain subregions. Furthermore, spatiotemporal analyses reveal that pseudogenes in different brain subregions have greatly varying temporal trajectories (e.g., increasing vs. decreasing), in contrast to protein-coding genes that tend to be more uniform. Finally, we identify a set of pseudogenes exhibiting significant changes in neuropsychiatric disorders, some of which overlap with known brain eGenes (genes whose expression is associated with expression quantitative trait loci, eQTLs), as well as with genes implicated by genome-wide association studies (GWAS). Together, our study provides a public resource of pseudogene expression in human and macaque brains (in comparison to other tissues) and highlights aspects of pseudogene expression in brain development and pathogenesis.

## Introduction

Pseudogenes are DNA sequences similar to protein-coding genes that contain disabling mutations rendering them unable to produce fully functional proteins^2^. Pseudogenes are categorized into three biotypes based on their mode of origin. Processed pseudogenes arise from retrotransposition events; duplicated pseudogenes, also referred to as unprocessed pseudogenes, originate from gene duplication events; unitary pseudogenes result from fixed loss-of-function (LoF) mutations in functional genes^3^. Processed and duplicated pseudogenes retain homologous protein-coding counterparts (i.e., parent protein-coding genes) within the genome, whereas unitary pseudogenes are defined by the presence of functional homologs in syntenic regions of other species. The study of pseudogenes presents technical challenges due to their high sequence similarity to parent protein-coding genes, necessitating high-quality genome annotation and accurate read alignments to distinguish pseudogenes from functional homologs^4^.

Traditionally, pseudogenes have been considered non-functional genomic elements that evolve under neutral selection, permitting the accumulation of deleterious mutations such as in-frame stop codons and frameshift insertions or deletions^5^. However, despite their characterization as “genomic fossils,” the last decade has witnessed numerous studies reporting that some pseudogenes are actively transcribed^6–10^. Specifically, they have been shown to regulate the expression of their functional protein-coding homologs by serving as a source of small interfering RNAs (siRNAs)^11^, microRNA binding sites^12^, or competing endogenous RNAs (ceRNAs)^13^.

Developmentally regulated pseudogenes have been catalogued across various human tissues^14^, and growing evidence links them to a range of diseases^15–17^—most notably cancer^18^. In the brain, where the transcriptome is exceptionally dynamic and tissue-specific, pseudogenes transcription is likewise pervasive^10^. Emerging work implicated pseudogenes transcribed in the brain in neurodegenerative disorders (for example, Alzheimer’s and Parkinson’s disease via retromer dysfunction^19^), psychiatric disorders such as autism spectrum disorder (ASD)^20^ and schizophrenia (SCZ)^21^, and intracranial malignancies such as glioma^22^. However, despite these individual findings, a systematic assessment of their collective role in major psychiatric disorders—ASD, SCZ, and bipolar disorder (BD)—has yet to be undertaken. As part of the PsychENCODE Project^1^, gene expression profiles have been characterized during different developmental stages and across psychiatric disorders using large-scale bulk RNA sequencing (RNA-seq) data from the human brain in several independent studies^23,24^. Despite these advances, pseudogenes remain understudied in these analyses due to their functional ambiguity and high sequence similarity to protein-coding genes, leading standard RNA-seq pipelines to frequently fail in accurately quantifying their expression. Therefore, a comprehensive analysis across large brain sample cohorts is essential to elucidate pseudogene expression dynamics in normal brain development and psychiatric disorders.

In this study, we aim to explore whether pseudogenes, traditionally considered as a group of non-functional remnants of protein-coding genes, could show patterns suggestive of putative roles in the brain. To that end, we propose three primary objectives: (i) characterizing the tissue- and brain region-specificity of pseudogene expression; (ii) identifying temporally dynamic (TD) pseudogenes across diverse brain regions; and (iii) identifying differentially expressed pseudogenes associated with psychiatric disorders. To achieve these aims, we integrated large-scale RNA-seq datasets from human and macaque brains, as well as human non-brain tissues. Leveraging publicly available RNA-seq data from the PsychENCODE Consortium^23,24^ and ENCODE EN-TEx project^25^, we applied a computational framework to accurately estimate pseudogene expression across a large number of human brain samples and ∼30 non-brain tissues. We then systematically characterized pseudogene expression specificity in the brain and examined their spatiotemporal dynamics during human brain development, comparing these patterns with those observed in the macaque brain. Finally, we identified a catalog of differentially expressed pseudogenes in psychiatric disorders (i.e., ASD, BD, and SCZ), revealing disorder-specific expression patterns. Taken together, this resource provides valuable insights into potentially functional pseudogenes in the human brain, advancing our understanding of their roles in neurodevelopment and psychiatric disorders.

## Results

### Accurate quantification of pseudogene expression in the human and macaque brains

It is challenging to accurately estimate expression levels of pseudogenes using short-read RNA-seq data due to the high sequence similarity between the pseudogenes and their parent protein-coding genes. To this end, we developed a computational framework aiming to detect actively transcribed pseudogenes and accurately quantify their expression levels and applied the computational framework to the RNA-seq datasets from human and macaque (**Figure 1A** and **1B**; **Supplementary Tables S1-S4**; see **Methods**). Briefly, we began with the pseudogene annotation from GENCODE^26^, and removed their exons overlapping with well annotated protein-coding genes and long non-coding RNAs (lncRNAs) at the gene level. We then retained and used the exons located in the uniquely mappable genomic regions for RNA-seq reads alignment. This approach enabled us to quantify pseudogene expression based on the RNA-seq reads aligned to the exons with high genome-wide sequence uniqueness, which could minimize ambiguous short read alignments. In addition, we excluded short exons to enhance specificity of short read alignments. As a result, we retained a set of 4,067 pseudogenes in the human genome that could be reliably quantified for their expression levels. For a fair comparison, we also applied the same framework to protein-coding genes in the human genome, yielding a set of 17,599 protein-coding genes in the downstream analysis.

**Figure 1.**
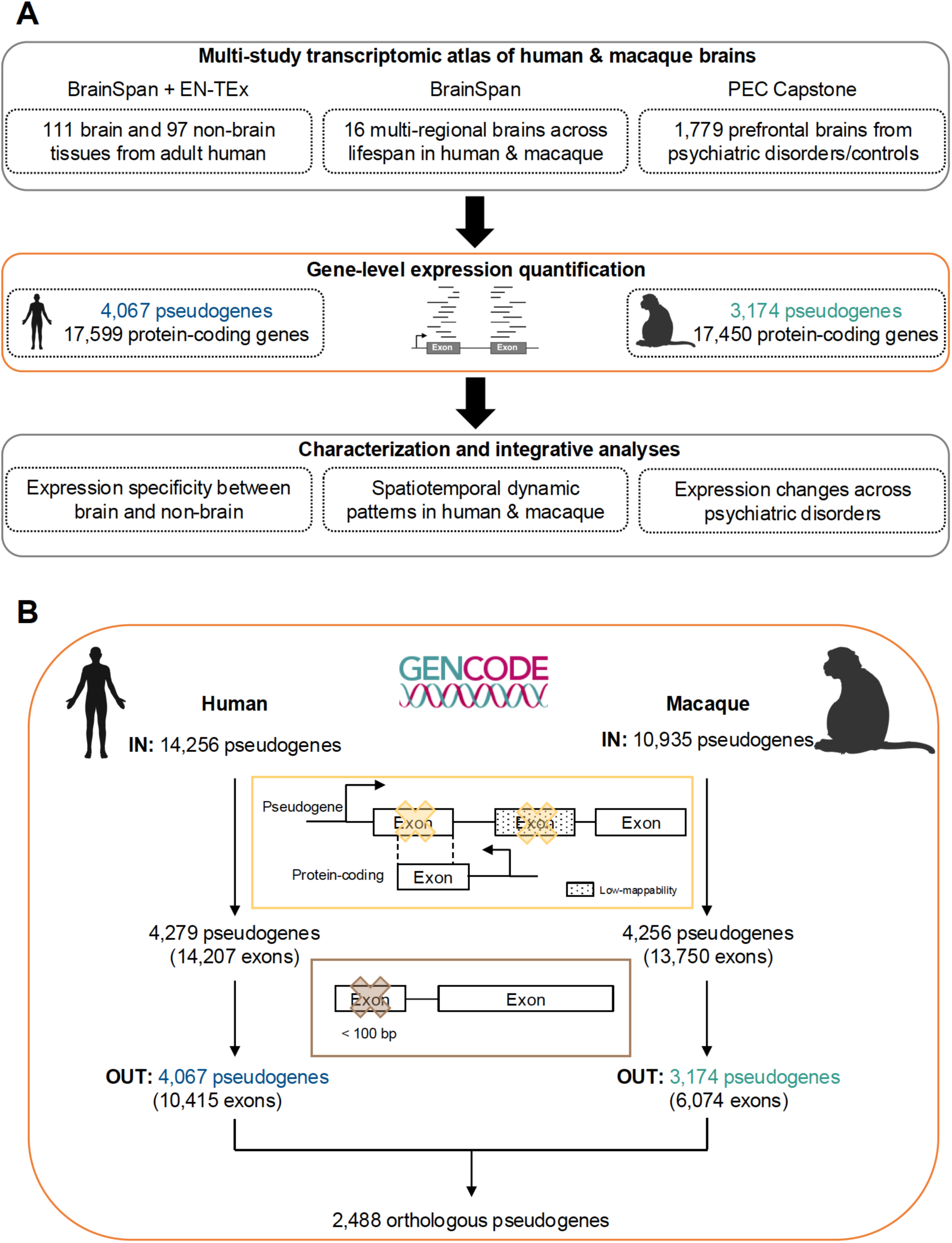
An overview of the analysis framework of this study. (**A**) Compilation of RNA-seq datasets from human and macaque, including: (i) non-brain tissues from human^25^; (ii) multi-regional brains from human and macaque^23^; and (iii) human prefrontal brains diagnosed with ASD, BD, SCZ, and neurotypical controls^24^. Notably, only the exons located in the genomic regions with high sequence uniqueness were used in the gene expression quantification, aiming to reduce read alignment ambiguity between pseudogenes and their parent protein-coding genes. The gene-level expression quantification was performed on the human and macaque reference genome, respectively. (**B**) Generating pseudogene annotations used in this study. Pseudogene annotations from GENCODE v39 were converted from hg38 to hg19 (human) and rheMac10 (macaque). Only the pseudogene exons retained after several filtering steps were used for the gene expression quantification. For macaque, only the duplicated and processed pseudogenes overlapping with PseudoPipe^27^ prediction were retained in the analysis. Panels (**A**) and (**B**) were partially created with BioRender.com.

Given the lack of high-quality pseudogene annotation for macaque genome, we first generated pseudogene annotation with high confidence by mapping the pseudogene annotation from the human genome onto the macaque genome (**Figure 1B**; see **Methods**). After this step, we had 10,935 pseudogenes (with 28,943 exons) annotated in the macaque genome. We then applied the same computational framework to the pseudogene annotation in the macaque genome. To improve the quality of pseudogene annotation in macaque, we also integrated the pseudogene annotation generated by PseudoPipe^27^ (**Supplementary Table S5**), resulting in a set of 3,174 pseudogenes in the macaque genome. Among these pseudogenes, 2,488 have definite orthologous relationships to the pseudogenes from human.

We then applied the computational framework to the RNA-seq datasets from diverse non-brain tissues in human collected from ENCODE EN-TEx project^25^. As expected, we confirmed distinct sample clustering in terms of their tissue source based on both pseudogenes and protein-coding genes (**Supplementary Figure S1**; **Supplementary Tables S6-S9**), indicating that the expression levels of pseudogenes estimated by our framework can represent the tissue identity as protein-coding genes. We also identified 5,108 pseudogene–parent gene pairs in the human genome, comprising 1,287 duplicated and 3,821 processed pseudogenes. Consistent with the previous study^7^, the duplicated pseudogenes were more likely to be located on the same chromosome as their parent protein-coding genes (57.3%) compared to the processed pseudogenes (4.9%). In addition, the genomic distance between duplicated (n = 357, adjusted R^2^ = 0.34, *P*-value < 2.2×10^-16^, F-test) and processed (n = 35, adjusted R^2^ = 0.46, *P*-value = 4.2×10^-6^, F-test) pseudogenes and their parent protein-coding genes showed significantly negative correlations with the genomic contact strength (**Supplementary Figure S2A**). The co-expression between duplicated pseudogenes and their parent protein-coding genes exhibited a strong positive correlation with their genomic proximity (**Supplementary Figure S2B**). Specifically, stronger expression correlation was observed when the duplicated pseudogenes and their parent protein-coding genes were located on the same chromosome and within 1 Mbp. In contrast, the processed pseudogenes did not show a similar correlation pattern as we observed in the duplicated pseudogenes. Taken together, these results suggest that our computational framework was effective for distinguishing the expression pattern of pseudogenes from that of their parent protein-coding genes despite their high sequence similarity, and the inferred expression levels are biologically meaningful and reliable for the downstream analysis.

### Strong expression specificity of pseudogenes across diverse tissue types and brain regions

Gene expression profiles from whole-body tissues and various brain regions in the adult human were analyzed to evaluate the tissue- and region-specificity of pseudogenes, their parent protein-coding genes, as well as total protein-coding genes. This analysis included 29 non-brain tissues (**Supplementary Tables S6** and **S7**), as well as 16 distinct brain regions across major anatomical structures (including the frontal lobe, temporal lobe, parietal lobe, occipital lobe, hippocampus, thalamus, basal ganglia, and cerebellum) (**Supplementary Table S8** and **S9**). For tissue-specificity, we considered genes that were transcribed in both brain and non-brain tissues, while for region-specificity, we focused exclusively on the genes transcribed within the brain (see **Methods**).

Across all the brain and non-brain samples, pseudogenes exhibited stronger expression specificity compared to their parent protein-coding genes and total protein-coding genes (**Figure 2A**). Notably, a significantly large proportion of pseudogenes (56.8%, 129 out of 227) displayed elevated expression in brain samples compared to their parent protein-coding genes (27.5%, 286 out of 1,041), and total protein-coding genes (29.2%, 3,570 out of 12,207) (**Figure 2B**). A large majority of pseudogenes with elevated expression in brain samples are processed pseudogenes (**Supplementary Figure S3**). Previous studies have revealed regulatory associations between processed pseudogenes and long interspersed elements (LINEs) in human genome^28,29^. Taken together, our findings confirm that processed pseudogenes and LINE-1 retrotransposons might be co-regulated and were specifically active in the human brain^30^.

**Figure 2.**
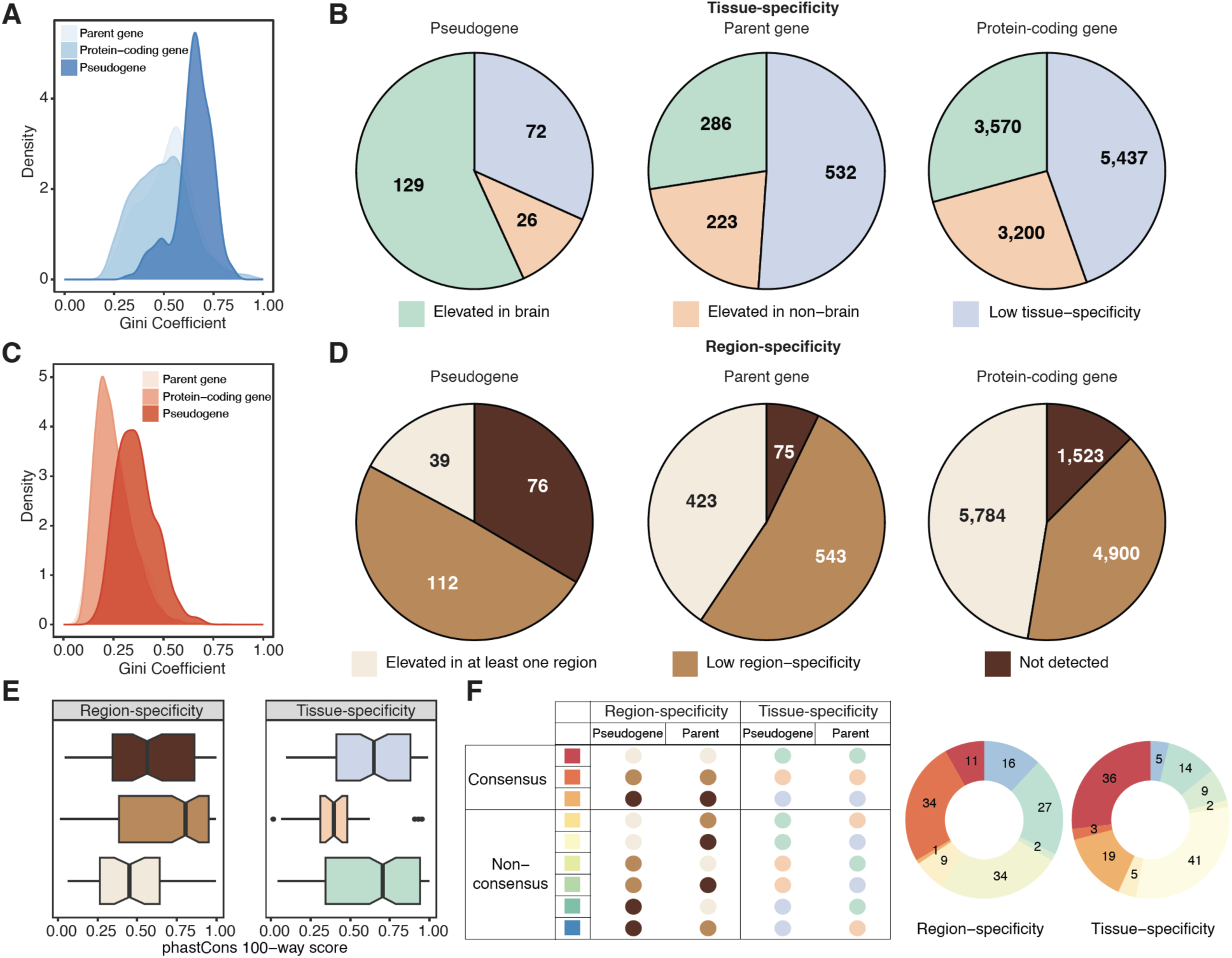
Tissue- and region-specificity of expression of pseudogenes, parent protein-coding genes, and protein-coding genes. (**A**) Distribution of the Gini coefficients for the expression of pseudogenes, parent protein-coding genes, and protein-coding genes across brain and non-brain tissues. (**B**) Number of differentially expressed genes (DEGs) among pseudogenes (left), parent genes (middle), and protein-coding genes (right) across brain and non-brain tissues. (**C**) Distribution of the Gini coefficient for the expression of pseudogenes, parent genes, and protein-coding genes across multiple brain regions. (**D**) Number of DEGs among pseudogenes (left), parent genes (middle), and protein-coding genes (right) across multiple brain regions. (**E**) Box plots showing the distribution of phastCons 100-way conservation scores of region- and tissue-specific pseudogenes. (**F**) Correspondence patterns between the pseudogenes and their parent genes with respect to region- and tissue-specificity. Pseudogene–parent gene pairs categorized into the same specificity class are defined as consensus pairs, whereas those falling into different classes are designated as non-consensus pairs.

Given that our gene expression profiles covered diverse brain regions from adult human, we were able to systematically evaluate the region-specificity of pseudogenes and their parent protein-coding genes in adult brain. Consistent with our findings on tissue-specificity, across brain regions, pseudogenes exhibited a stronger expression specificity compared to their parent protein-coding genes and total protein-coding genes (**Figure 2C**). Notably, a larger proportion of actively transcribed pseudogenes (49.3%, 112 out of 227) were broadly expressed across diverse brain regions compared to total protein-coding genes (40.1%, 4,900 out of 12,207) (**Figure 2D**). As expected, we found that protein-coding genes with elevated expression in brain were significantly enriched in synaptic signaling and brain disorder-associated biological pathways. Additionally, the protein-coding genes with elevated expression in at least one brain region were enriched in the biological functions related to neurogenesis and signaling pathways (**Supplementary Figure S4**).

We systematically compared the evolutionary conservation and population variation of pseudogenes with tissue- and region-specific expression patterns to better understand the potential functions of those with elevated expression in the brain or broadly expressed across brain regions. We found that the pseudogenes with elevated expression in the brain and broad expression across different brain regions were more conserved than their companion sets. However, no significant difference was observed in the proportion of rare variants (**Figure 2E**; **Supplementary Figures S5** and **S6**). These results suggest that while some pseudogenes in the human brain are under functional constraints (i.e., more evolutionarily conserved), they are not subject to strong purifying or positive selection to the same extent as protein-coding genes.

### Concordance of expression specificity between pseudogenes and their parent protein-coding genes

We investigated the expression specificity of pseudogenes and their parent protein-coding genes across human tissues and brain regions. We observed that pseudogenes with elevated expression in the brain tend to show broader expression across diverse brain regions compared to protein-coding genes (**Supplementary Figure S7**). In previous results, we showed that the pseudogenes and their parent protein-coding genes may have co-expression due to 3D chromatin architecture. Thus, we then explored the expression concordance between pseudogenes and their parent protein-coding genes (**Figure 2F**). Overall, more pseudogene–parent gene pairs showed concordant expression at the tissue level than at the brain region level. Interestingly, many pseudogenes displayed strong tissue-specific expression with elevated levels in the brain, while their parent protein-coding genes were more broadly expressed across tissues. Conversely, within the brain, pseudogenes tended to be expressed across multiple regions, whereas their parent genes showed more regionally restricted expression. These observations suggest that pseudogenes and their parent protein-coding genes display distinct, yet partially overlapping, expression patterns across diverse tissues and brain regions.

### Spatiotemporal dynamics of pseudogene expression in human and macaque brain

Previous studies have revealed that human brain transcriptome undergoes pervasive spatiotemporal dynamic changes, particularly during the prenatal stages^31–33^. However, the spatiotemporal expression dynamics of pseudogenes have not been explored, and even less is known about their spatiotemporal expression dynamics in macaque brain. To this end, we leveraged a set of RNA-seq datasets from 603 human (**Supplementary Tables S8** and **S9**) and 430 macaque (**Supplementary Tables S10** and **S11**) brains across lifespan—covering 16 different brain regions—to detect the pseudogenes alongside protein-coding genes exhibiting temporally dynamic (TD) patterns during brain development and aging. Briefly, we characterized four distinct TD gene expression patterns—rising, falling, U-shaped, and inverted U-shaped—in both human and macaque brains and assigned each TD gene to one of these categories (**Supplementary Tables S12** and **S13**; see **Methods**). We then compared the proportions of these four trajectory categories between pseudogenes and protein-coding genes across brain regions in both species (**Figure 3A** and **3B**; **Supplementary Figure S8**).

**Figure 3.**
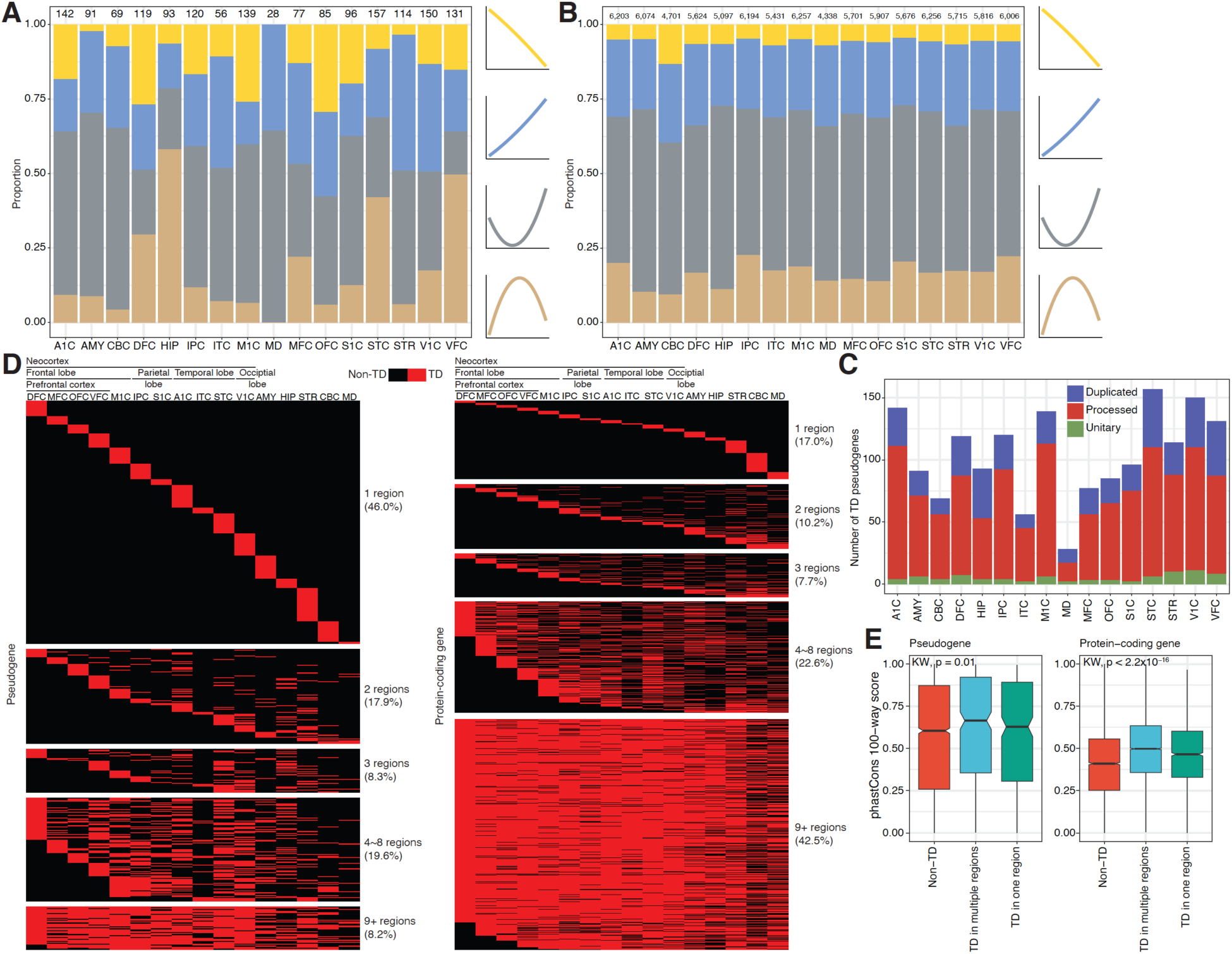
Spatially and temporally dynamic expression of pseudogenes and protein-coding genes in the human brains. (**A**) Proportion of different classes of temporally dynamic pseudogenes across multiple brain regions. (**B**) Proportion of different classes of temporally dynamic protein-coding genes across multiple brain regions. (**C**) Number of temporally dynamic pseudogenes, categorized by type, across multiple brain regions. (**D**) Heatmaps showing temporally dynamic pseudogenes (left) and protein-coding genes (right) that are uniquely identified in a single brain region or shared across multiple regions. (**E**) Box plots showing the distribution of phastCons 100-way conservation scores of temporally dynamic pseudogenes (left) and protein-coding genes (right). KW, Kruskal-Wallis; TD, temporally dynamic.

The total number of TD protein-coding genes in human brains varied in terms of region. However, the relative proportions of different types of TD protein-coding genes (i.e., rising, falling, U-shaped, and inverted U-shaped) were generally stable, with the U-shaped ones as the most common type. We observed fewer TD pseudogenes in macaque brains compared to human brains. The TD pseudogenes exhibited stronger variety in terms of their TD types than the protein-coding genes. Most of the TD pseudogenes were processed pseudogenes (68.8% in human and 69.2% in macaque) (**Figure 3C**; **Supplementary Figure S9**). These results revealed that the pseudogenes—especially the processed ones—exhibited more complex spatiotemporal expression patterns.

Having established overall spatiotemporal trends, we next explored the region-specific characteristics of these TD genes. In human, we found that a substantial proportion of the TD pseudogenes exhibited region specificity, with 46.0% identified as TD in a single brain region. In contrast, TD protein-coding genes displayed a broader distribution, as only 17% are TD in single brain region, while the majority (65.1%) were shared across multiple regions (**Figure 3D**). Similarly, in macaque, 58.8% of the TD pseudogenes were restricted to a single brain region, whereas the majority (52.9%) of the TD protein-coding genes were shared across multiple regions (**Supplementary Figure S10**).

We further compared sequence conservation for genes identified as TD versus non-TD, as well as the TD genes restricted to single or multiple brain regions. For both pseudogenes and protein-coding genes, we detected significant overall differences in conservation (*P*-value = 0.01 for pseudogenes, and *P*-value < 2.2×10^-16^ for protein-coding genes, Kruskal–Wallis test) (**Figure 3E**). However, the proportion of rare variants in pseudogenes did not differ among these groups (**Supplementary Figure S11**). These results suggest that the TD pseudogenes exhibited a notable degree of sequence conservation, indicating potential functionality in brain development. The TD genes expressed in multiple brain regions appear to be the most conserved, implying that they were subject to stronger selective pressures. Nevertheless, as expected, in comparison to the TD protein-coding genes, pseudogenes seemed to experience weaker evolutionary constraints, which may explain why there was no substantial reduction in their rare variant frequency.

### Inferring brain-associated functions of pseudogenes

The biological functions of pseudogenes are poorly explored, as they are frequently presumed to lack functions^4^. It will be useful to investigate the putative brain-associated biological processes in which the pseudogenes may be implicated by leveraging the expression profiles of pseudogenes and protein-coding genes estimated based on the computational framework in this study.

Although we have identified the brain regions and developmental stages in which specific pseudogenes exhibit elevated expression, we cannot clearly predict their biological functions, because every brain region or developmental stage has multiple enriched biological functions, and we cannot assign the associated functions to specific pseudogenes. Instead, we hypothesize that the pseudogenes in co-expression with protein-coding genes within the same modules may participate in similar biological processes. To this end, we constructed a set of co-expression modules of pseudogenes and protein-coding genes to explore the enriched biological functions of the pseudogenes using weighed gene co-expression network analysis (WGCNA). We analyzed a set of 927 expressed pseudogenes and 13,340 expressed protein-coding genes in human, as well as 275 expressed pseudogenes and 13,165 expressed protein-coding genes in macaque. Finally, an average of 20 and 14 co-expression modules across 16 brain regions were obtained in human and macaque, respectively. We observed substantial regional variability in module sizes across human and macaque brain regions. For instance, in the human brain, the DFC and CBC exhibited large median module sizes (230 and 225 genes, respectively), whereas the HIP and M1C showed much smaller medians (108 and 111 genes, respectively). This variation likely reflects differences in the complexity of transcriptional architecture across brain regions. (**Figure 4A**; **Supplementary Figure S12A**). In total, we found that 704 pseudogenes (75.9%) in human and 197 pseudogenes (71.6%) in macaque exhibit obvious co-expression with the protein-coding genes. The proportion of pseudogenes in co-expression modules is markedly lower in macaque compared to those in human (**Figure 4B**; **Supplementary Figure S12B**). Although the majority of modules are predominantly composed of protein-coding genes, a distinct subset demonstrates a significant enrichment for pseudogenes, particularly in the human brain.

**Figure 4.**
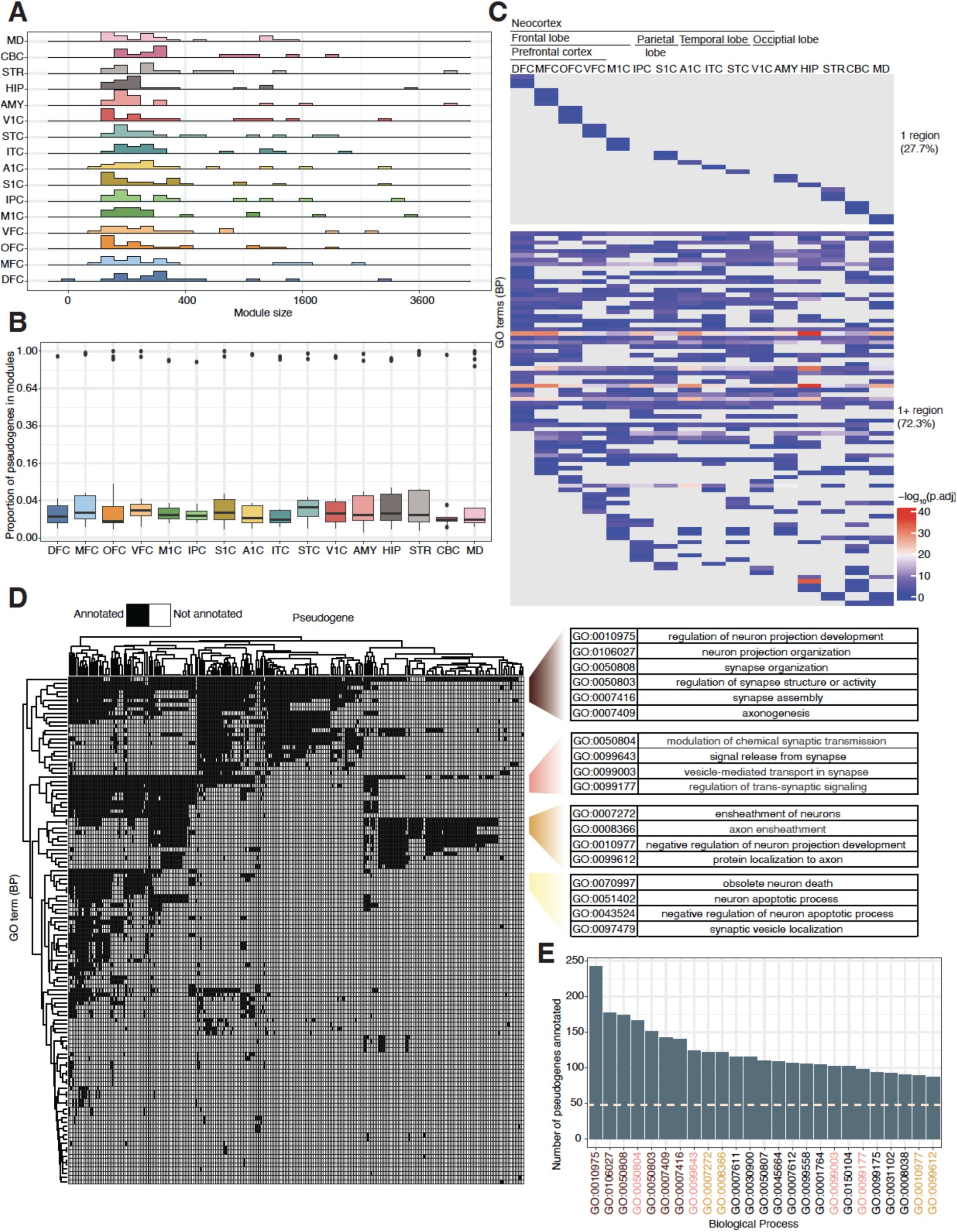
Functional inference of pseudogenes using WGCNA in the human brain. (**A**) Ridge plot showing the distribution of co-expression module sizes (excluding Module 0) across different brain regions. (**B**) Box plot showing the proportion of pseudogenes within co-expression modules (excluding Module 0) for each brain region. (**C**) Heatmap showing enriched brain function-associated Gene Ontology (GO) terms derived from co-expression modules containing pseudogenes, categorized as either region-specific or shared across multiple brain regions. (**D**) Heatmap showing the GO terms annotated for individual pseudogenes. Selected representative GO terms and their functional descriptions are listed. (**E**) Number of pseudogenes associated with each GO term. Only the top 25 terms are shown. The dashed horizontal line indicates the average number of pseudogenes for each GO term. BP, biological process.

We then performed Gene Ontology (GO) enrichment analysis on the genes within each module to infer their potential biological functions, focusing on terms associated with brain or neuronal processes. Specially, we identified several GO terms with strong relevance to neural activity and brain development. For instance, regulation of neuron projection development (GO:0010975) and axonogenesis (GO:0007409) are fundamental neurodevelopmental processes that contribute to the establishment of neuronal polarity, axonal outgrowth, and the formation of functional neural circuits^34^. These terms reflect essential biological processes involved in neural communication, circuit formation, and signal transduction, underscoring the potential roles of these co-expressed genes in brain-specific functions. Also, 27.7% of the GO terms were unique to a single brain region, whereas the remainder were shared across multiple regions (**Figure 4C**). This finding highlights the region-specific functional organization of gene co-expression modules, suggesting that distinct brain regions may harbor specialized transcription programs contributing to neurodevelopmental and cognitive regulations. These observations are consistent with prior transcriptomic studies demonstrating spatial specificity in gene expression across brain regions^31^. Next, we inferred the putative functions for the pseudogenes in the co-expression modules by assigning the significantly enriched GO terms to them. Functional annotations derived from different brain regions were aggregated to generate a comprehensive catalog of GO terms associated with each pseudogene (**Figure 4D**). Notably, we observed that individual GO terms were frequently linked to a substantial number of pseudogenes, with a median of 47.2 pseudogenes annotated for each GO term (**Figure 4E**).

### Shared pseudogene dysregulation across psychiatric disorders

Large-scale initiatives such as ROSMAP^35^ and PsychENCODE^36^ have significantly advanced our understanding on the gene expression in psychiatric and degenerative disorders. However, the expression patterns of pseudogenes in these brain disorders are poorly characterized. Here, we aim to fill this gap by analyzing the differential expression of pseudogenes in psychiatric disorders, including ASD, BD and SCZ (**Supplementary Tables S14** and **S15**; see **Methods**).

We identified a set of differentially expressed pseudogenes and protein-coding genes in each of the psychiatric disorders (**Supplementary Table S16**). Compared to protein-coding genes, pseudogenes exhibited significantly greater expression fold changes (*P*-value = 3.97×10^-8^ for downregulated and *P*-value = 1.74×10^-23^ for upregulated genes in ASD; *P*-value = 1.16×10^-19^ for downregulated and *P*-value = 3.52×10^-8^ for upregulated genes in BD; *P*-value = 6.24×10^-50^ for downregulated and *P*-value = 1.02×10^-16^ for upregulated genes in SCZ, FDR-adjusted, Wilcoxon rank sum test) (**Figure 5A**), suggesting that these pseudogenes may not be non-functional elements, but instead possess putative regulatory roles in the context of psychiatric disorders. Furthermore, similar to protein-coding genes, we observed a considerable proportion (22.1% of the upregulated and 26.5% of the downregulated) of the differentially expressed pseudogenes that were shared among ASD, BD and SCZ (**Figure 5B**; **Supplementary Figure S13**). Differentially expressed protein-coding genes across the psychiatric disorders were enriched in the biological processes related to immune response, synaptic signaling, cell cycle regulation, and neurodevelopment, with both shared and disorder-specific patterns observed among ASD, BD and SCZ (**Supplementary Figure S14**). In parallel, cell type enrichment analysis revealed significant associations between dysregulated genes and diverse brain- and immune-related cell types (**Supplementary Figure S15**). Previous studies have revealed the convergent molecular mechanisms of these different psychiatric disorders by analyzing the overlap of differentially expressed protein-coding genes^37–39^. These results indicate that pseudogenes may be implicated in the convergent molecular mechanisms of different psychiatric disorders.

**Figure 5.**
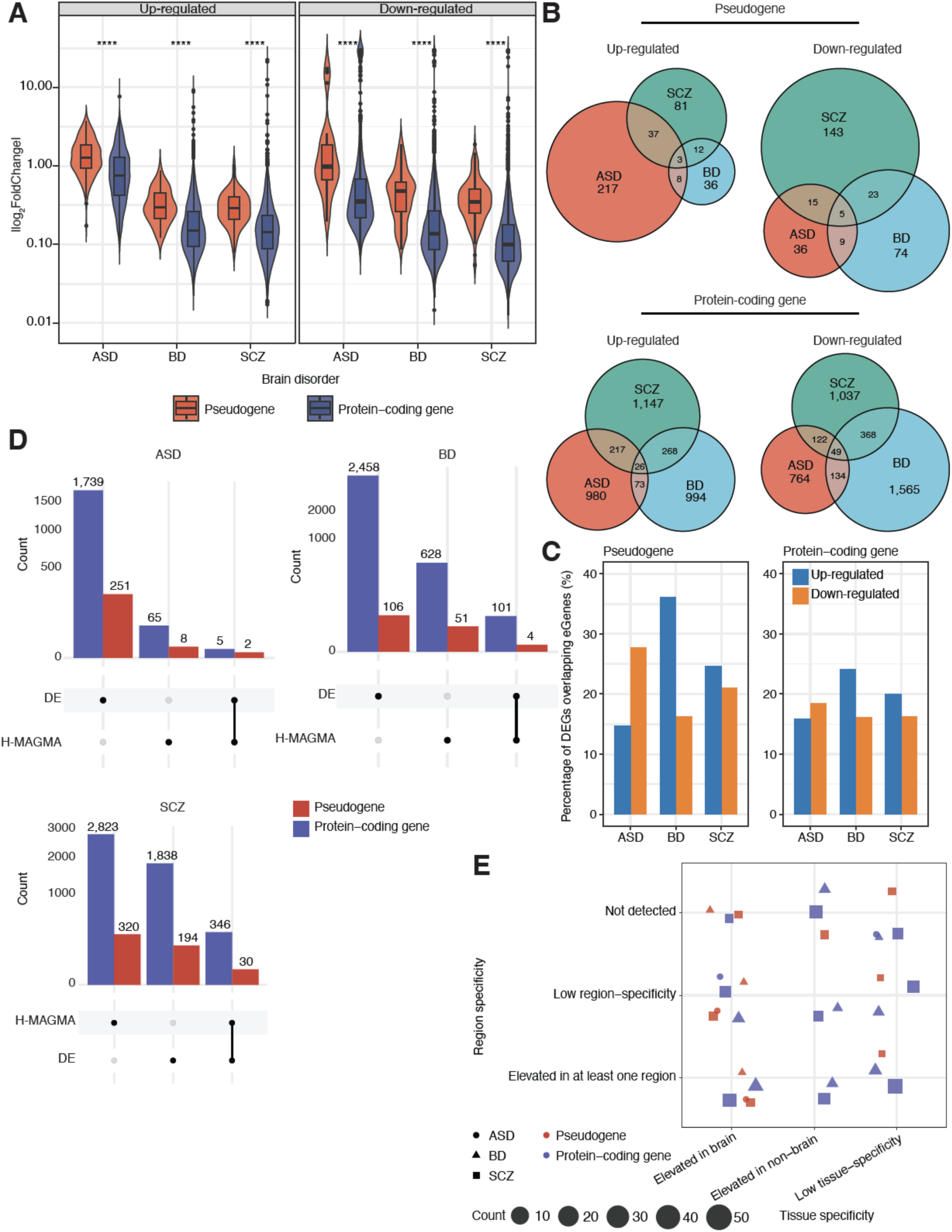
Differential expression of pseudogenes and protein-coding genes across psychiatric disorders. (**A**) Violin plots showing the fold changes of upregulated (left) and downregulated (right) protein-coding genes and pseudogenes identified in autism spectrum disorder (ASD), bipolar disorder (BD), and schizophrenia (SCZ). ****, *P*-value < 0.0001. (**B**) The number of upregulated and downregulated protein-coding genes (top) and pseudogenes (bottom) shared among ASD, BD, and SCZ. (**C**) Bar plots showing the percentage of differentially expressed genes that overlap with eGenes derived from GTEx v7 (hg19 based), shown separately for pseudogenes (left) and protein-coding genes (right). (**D**) Overlap between the DEGs and the disorder risk genes prioritized by H-MAGMA^40^, shown for both pseudogenes and protein-coding genes. (**E**) Region- and tissue-specificity of the overlapping genes identified in (**D**), shown for both pseudogenes and protein-coding genes.

To further elucidate the regulatory logics underlying the expression changes of these differentially expressed pseudogenes, we performed integrative analysis for these pseudogenes and expression quantitative trait loci (eQTL). Similar to protein-coding genes, a subset of differentially expressed pseudogenes was found to be eQTL-linked (i.e., eGenes), suggesting that specific single nucleotide polymorphisms (SNPs) may regulate the expression of these pseudogenes (**Figure 5C**). More importantly, the disorder risk genes prioritized by H-MAGMA^40^ (**Supplementary Table S17**) showed overlap with the differentially expressed pseudogenes in ASD (2 pseudogenes), BD (4 pseudogenes), and SCZ (30 pseudogenes) (**Figure 5D**). Notably, SCZ exhibited a higher number of overlapping genes compared to ASD and BD. This convergence between differentially expressed pseudogenes and H-MAGMA-derived risk loci implies that their dysregulation may be driven by genetic variants implicated in disease susceptibility. To further characterize these pseudogenes shared between differential expression analyses and H-MAGMA risk loci, we investigated their region- and tissue-specific expression patterns (**Figure 5E**). We identified several SCZ-associated pseudogenes with elevated expression in the brain and evidence of region-specificity. Collectively, these findings support the notion that pseudogenes may have functional relevance beyond mere transcriptional artifacts, potentially mediating the effects of inherited genetic risk in psychiatric conditions.

## Discussion

The functionality of specific pseudogenes has been known for decades^4,41–44^. Yet, the expression patterns of pseudogenes in brain and their putative associations with psychiatric disorders are still not fully explored. In this study establishing a computational framework capable of minimizing the expression bias derived from their high sequence similarity to their parent protein-coding genes, we analyzed the pseudogene expression profiles based on large-scale RNA-seq data from the human and macaque brains. We characterized a set of actively transcribed pseudogenes in the brains, which showed strong tissue and region specificity, as well as distinct temporal dynamic patterns during brain development. We also identified differentially expressed pseudogenes in different psychiatric disorders, and revealed the regulatory logics of these pseudogenes, indicating their implications in the pathogenesis of psychiatric disorders.

Although pseudogene expression profiles in human brain have been recently generated^24^, the computational framework for gene expression quantification was not developed by carefully considering the scenario of high sequence similarity between pseudogenes and their parent protein-coding genes. Furthermore, accurate quantification of pseudogene expression and analyses of their expression patterns in non-human primates are still lacking, which may hinder the exploration on the biological functions of pseudogenes. Pseudogenes have been revealed as new players in complex human diseases^15,45–47^. Some transcribed pseudogenes share miRNA binding sites with their parent protein-coding genes, thus enabling competing for the same miRNAs in the context of cancer and neuron^12,48–50^. However, the expression changes of pseudogenes in psychiatric disorders have not been well characterized, and the regulatory mechanisms of pseudogenes in psychiatric disorders are still unclear.

One limitation of this study is that, despite restricting our analysis to uniquely mappable genomic regions, short-read RNA-seq reads can still map equally well to both pseudogenes and their corresponding parent protein-coding genes, which could potentially lead to bias expression quantification. Furthermore, the exclusion of certain exonic regions may contribute to the underestimation of gene expression. With the increasing availability of long-read RNA-seq datasets, such as those generated through the capture long-read sequencing strategy^51^, it is now possible to achieve greater sensitivity in detecting expression changes, particularly in pseudogenes. This is also exemplified by recent research focusing on the *GBA1* locus, where long-read RNA-seq enabled clear resolution of *GBA1* and its pseudogene *GBAP1* in the human brain^52^. In addition, the recent Telomere-to-Telomere (T2T) projects for non-human primates (NHPs) has produced a complete genome assembly of the macaque^53^, facilitating more accurate genome annotation and transcriptomic analyses. Complementing these advances, the developmental Genotype-Tissue Expression (dGTEx) project offers extensive transcriptomic data across multiple developmental stages in both humans and NHPs^54^, enabling novel insights into the regulatory mechanisms of pseudogene expression during development.

In summary, our findings suggest that pseudogenes are implicated in brain function and the pathophysiology of psychiatric disorders, challenging the conventional view of pseudogenes as mere genomic fossils. This work lays a foundation for future investigations into the mechanistic roles of pseudogenes in the brain, opening new avenues for understanding complex psychiatric disorders from a non-coding genome perspective.

## Methods

### Generating pseudogene annotations in the human and macaque genomes

All brain and non-brain tissue BAM files used in this study were aligned to the GRCh37 assembly. Accordingly, we used the GRCh37.p13 reference genome assembly and the GENCODE v39 genome annotation^26^ that was mapped back to the GRCh37 assembly when uniformly processing the large-scale BAM files from the human brain and non-brain tissues. For macaque, we used the most up-to-date rheMac10 reference genome (released in February 2019), given the substantial improvements of genome sequencing quality for this non-human primate. The macaque reference annotation was derived by mapping the human reference annotation to the macaque genome using Liftoff^55^, with the options -exclude_partial and -polish. As a result, 10,935 pseudogenes (76.7% of 14,256) and 19,131 protein-coding genes (95.1% of 20,111) from the human genome were successfully mapped to the macaque genome.

We computed the mappability scores for the human and macaque genomes using GEM^56^ (version 2.0.14.1) with a window size of 100 base pairs to avoid confounding from ambiguous read alignments. To accurately identify pseudogenes, we extracted the exons from genome annotations and excluded those exons overlapping with annotated protein-coding genes and lncRNAs. Only exons located within uniquely mappable genomic regions (mappability scores > 0.95) were retained. Next, the exons shorter than 100 base pairs were removed. In total, we retained 4,067 pseudogenes in human for the downstream analysis.

For the pseudogenes in human genome, we directly obtained their types (i.e., processed, duplicated, or unitary) from GENCODE v39 genome annotation^26^ that was mapped back to the GRCh37 assembly. For the pseudogenes in macaque genome, their genomic loci and types were determined using Liftoff^55^ from the human to the macaque genome. To enhance specificity, we further selected duplicated and processed pseudogenes by comparing their exons with annotations from PseudoPipe^27^, which was executed using default parameters. We employed BEDTools^57^ (version 2.30.0) to estimate the regional overlap, and retained the exons for which at least 50% of their length overlapped with the PseudoPipe annotations. In total, we retained 3,174 pseudogenes in macaque for the downstream analysis.

### Quantifying expression of pseudogenes in human and macaque

Leveraging multiple publicly available transcriptomic datasets, we quantified pseudogene and protein-coding gene expression in human brain and non-brain tissues as well as macaque brain tissues. We retained the samples with RNA integrity numbers (RINs) larger than 5.5 in the downstream analysis. Consequently, our study included 603 human brain samples spanning various developmental stages, as well as 1,779 human brain samples from the individuals diagnosed with autism spectrum disorder (ASD), bipolar disorder (BD), and schizophrenia (SCZ), alongside neurotypical controls obtained from the PsychENCODE project^23,24^. We also incorporated 98 non-brain tissue samples from the ENCODE EN-TEx project^25^ in the analysis.

For human brain and non-brain samples, we quantified pseudogene and protein-coding gene expression from existing BAM files using Cufflinks^58^ (version 2.2.1), generating the expression levels in both Fragments Per Kilobase of transcript per Million mapped reads (FPKM) and raw counts. Gene expression quantification was performed with the supplied reference annotation, in which exons were filtered (-G), and strand-specific information was incorporated (–library-type). The resulting expression estimates were merged to generate expression matrices for pseudogenes and protein-coding genes across three datasets. For macaque brain samples, since the original reads were mapped to the rheMac8, we first extracted FASTQ records using BEDTools bamtofastq^57^ and subsequently remapped the reads to the rheMac10 using STAR^59^ (version 2.7.9a). Expression quantification for macaque samples was performed with the BAM files using Cufflinks^58^, following the same approach for the human samples.

### Expression analysis between pseudogenes and their parent protein-coding genes

We used PseudoPipe^27^ to infer the parental relationships for the duplicated and processed pseudogenes in the GRCh38 assembly. Briefly, we intersected the results from PseudoPipe^27^ with the pseudogenes annotated in the Ensembl v103. The parental information of the pseudogenes in human genome was available in **Supplementary Table S18**. Finally, we converted these annotations back to the GRCh37, yielding a stable set of pseudogenes and their corresponding parent protein-coding genes.

To investigate how pseudogenes may interact with their corresponding parent protein-coding genes in the human brain, we analyzed Hi-C interaction maps from the human dorsolateral prefrontal cortex (DLPFC), obtained from the PsychENCODE portal (http://resource.psychencode.org/Datasets/Pipeline/HiC_matrices/PIP-01_DLPFC.10kb.txt.tar.gz). For pseudogenes and their parent protein-coding genes located on the same chromosome, we calculated their genomic distance and assigned Hi-C interaction counts based on the corresponding values in the matrix. Pseudogene–parent gene pairs located on different chromosomes were labeled as ‘NA.’ Pairs with a Hi-C count of zero were excluded from further analysis. We then performed a linear regression analysis by plotting the logarithm of the Hi-C count against the logarithm of the genomic distance (in kilobase pairs) between pseudogenes and their respective parent protein-coding genes.

Next, we examined the co-expression between the pseudogenes and their parent protein-coding genes. Given the distinct mechanisms of pseudogene biogenesis, we categorized pseudogene–parent gene pairs into three groups: (i) located on different chromosomes, (ii) located on the same chromosome with a genomic distance less than 1 Mb, and (iii) located on the same chromosome with a genomic distance greater than 1 Mb. The Pearson correlation coefficient was used to quantify co-expression for 5,108 pairs of pseudogenes and parent protein-coding genes across 1,779 brain samples from individuals with ASD, BD, SCZ, and neurotypical controls.

### Characterizing tissue- and region-specific expression patterns of pseudogenes

To ensure a robust comparison of pseudogene expression, we only selected adult samples (days > 6841) from the human brain dataset as all donors from the ENCODE EN-TEx project^25^ were adults. We identified 12,207 protein-coding genes as actively transcribed asking that the FPKM value of the gene was larger than 2 in at least one-third of the total samples (N = 208). Similarly, 619 pseudogenes were identified as actively transcribed based on the same thresholds, except that the threshold of FPKM values of the pseudogenes was 0.5 as pseudogenes generally showed lower expression than the protein-coding genes. Notably, we retained 1,477 pseudogenes that were located within the uniquely mappable regions and had reliably established parent protein-coding gene assignments in this analysis, aiming to make a fair comparison between the pseudogenes and their corresponding parent protein-coding genes.

To quantify the unevenness of gene expression across different samples, we calculated the Gini coefficients, which measure expression disparity^60^. The Gini coefficient is calculated as: 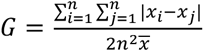, where *x_i_* and *x_j_* represent the expression level of a gene in samples *i* and *j*, respectively, *x̄* is the mean expression level across all samples, and *n* is the number of samples.

When studying tissue-specific gene expression, we applied DESeq2^61^ to analyze the expression of pseudogenes, their parent protein-coding genes, and total protein-coding genes using samples from the PsychENCODE Consortium^23^ and the ENCODE EN-TEx project^25^. Our objective was to identify the genes with elevated expression in either brain or non-brain tissues, as well as those exhibiting low tissue-specificity. A gene was considered tissue-specific if it met the criteria of |log2FoldChange| > 1 and an adjusted *P*-value < 0.05. For region-specific gene expression within the brain, we evaluated 120 unique pairwise comparisons across 16 distinct brain regions. We applied DESeq2^61^ to pseudogenes, their parent protein-coding genes, and total protein-coding genes using only samples from the PsychENCODE Consortium^23^. A gene was classified as region-specific if it met the criteria of |log2FoldChange| > 1 and an adjusted *P*-value < 0.05 in any of these pairwise comparisons.

Additionally, for region-specific gene expression within the brain, we considered only genes detected in the brain. As a result, we applied a criterion requiring at least 80% of samples to exhibit an FPKM > 0.5 for pseudogenes and FPKM > 2 for protein-coding genes. For those that are detected, we further consider whether they are region-specific or show a broad expression across regions.

### Sequence conservation analysis of pseudogenes

The 100-way PhastCons scores obtained from the UCSC Genome Browser were used to evaluate sequence conservation^62^. We then utilized bigwigAverageOverBed^63^ to compute conservation scores separately for exons of protein-coding genes and pseudogenes. To represent the overall sequence conservation level of each gene, we then calculated a weighted average across all exons.

To assess population-level variation, we obtained the genome-wide single nucleotide variation (SNV) data from gnomAD^64^ (v2.1.1), all of which were mapped to the GRCh37 assembly. The non-Finnish European (NFE) population was designated as the reference group. We retained only variants that passed quality control, exhibited an allele frequency (AF) of ≥ 0.001 in the NFE population, and had a prediction probability > 0.8 of being a true variant. Using BEDTools intersect^57^, we quantified the number of rare (AF ≤ 0.05) and common (AF > 0.05) variants within the gene body regions, and subsequently calculated the proportion of the rare variants.

### Temporal expression analysis of pseudogenes across different regions in human and macaque brains

We employed maSigPro^65^ to identify TD gene expression patterns across different regions of human and macaque brains. To account for potential non-linear relationships in gene expression during brain development, we set the model to fit a quadratic regression model. Specifically, genes with temporally dynamic expression changes were identified using a false discovery rate (FDR) threshold of 0.05, adjusted via the BH correction. Subsequently, a two-way backward stepwise selection procedure was implemented to refine the gene set by eliminating non-significant terms, and the final set of well-fitted genes was retained based on an R^2^ threshold of > 0.3. TD genes were categorized into four patterns: rising, falling, U-shaped (initial decrease followed by an increase), and inverted U-shaped (initial increase followed by a decrease), based on the observed time range.

### Co-expression analysis of pseudogenes and protein-coding genes in human and macaque brains

We utilized the WGNCA package^66^ to infer pseudogene functions across different human and macaque brain regions based on the large co-expression dataset. Initially, we filtered out low-expression genes, setting a threshold of FPKM ≥ 1 for protein-coding genes and FPKM ≥ 0.5 for pseudogenes in at least 200 samples out of 603 human samples. Similarly, we set the same threshold for pseudogenes and protein-coding in at least 100 samples out of 430 macaque samples. Next, we ran the goodSamplesGenes() function to further filter out genes and samples with too many missing entries. To construct a scale-free network for each region, we employed the pickSoftThreshold() function to determine the optimal soft-thresholding power. Subsequently, we used the blockwiseModules() function for automated network construction and module detection, enabling efficient analysis in a block-wise manner. The parameters for the function were set as follows: TOMType = ‘signed’, minModuleSize = 30, networkType = ‘signed’, mergeCutHeight = 0.25, detectCutHeight = 0.995, deepSplit = 2, minCoreKME = 0.7, minCoreKMESize=3, minKMEtoStay=0.7, pamRespectsDendro = TRUE. Finally, we saved the topological overlap matrix (TOM) for each human and macaque brain region for further analysis, including quantifying the number of pseudogenes in each module and performing GO enrichment analysis to infer pseudogene functions within the identified modules. For subsequent analyses, the grey module (Module 0) was excluded, as it comprises genes that do not exhibit strong co-expression relationships with other genes and therefore lack reliable network-based functional context.

We used clusterProfiler^67^ to perform both Gene Ontology (GO) and Kyoto Encyclopedia of Genes and Genomes (KEGG) pathway enrichment analysis for protein-coding genes with elevated expression in the brain, non-brain tissues, and at least one brain region, applying Benjamini-Hochberg (BH) correction for multiple comparisons with an adjusted *P*-value cutoff of 0.05, and extracting the top five terms based on the lowest adjusted *P*-values.

To ensure sufficient statistical power, only co-expression modules containing more than 30 genes were included. Enrichment significance was assessed using the BH method to correct for multiple testing, with an adjusted *P*-value threshold of 0.05. To focus on GO terms relevant to the brain- or neuron-related function, only the GO terms containing any of the following keywords—brain, neuron, axon, synaptic, synapse, learning, or memory—were retained for further analysis.

### Differential expression analysis in psychiatric disorders

To investigate the dysregulation of pseudogenes across psychiatric disorders, we performed pairwise differential gene expression analyses between the individuals diagnosed with specific psychiatric disorders and normal controls using DESeq2^61^. Multiple confounding factors from the metadata of samples—including sex, age at death, diagnosis, study, postmortem interval (PMI), RNA integrity number (RIN), library preparation protocol, strandedness, and sequencing platform—were regressed before the differential gene expression analyses. Surrogate variable analysis (SVA) was conducted using the R package sva^68^ to estimate five latent factors, which were then incorporated into the design model to account for hidden sources of variation. Genes were considered significantly differentially expressed based on an adjusted *P*-value threshold (e.g., FDR < 0.05).

To functionally characterize the differentially expressed protein-coding genes, we conducted GO and cell type enrichment analyses. For GO enrichment, upregulated and downregulated genes in each disorder were analyzed separately using the R package clusterProfiler^67^, applying the BH correction for multiple testing and a minimum gene set size threshold of three. The top seven enriched GO terms, ranked by adjusted *P*-values, were visualized for interpretation. For cell type enrichment, we obtained marker genes representing various human cell types from the CellMarker 2.0 database^69^ (http://bio-bigdata.hrbmu.edu.cn/CellMarker/). Similar to the GO enrichment analysis, cell type enrichment was conducted separately for upregulated and downregulated genes in each disorder, and the top seven enriched cell types were visualized based on adjusted *P*-values.

We leveraged the eGenes in the Brain Frontal Cortex (BA9) from GTEx^70^ v7 (hg19 based) data portal to explore which differentially expressed genes could be influenced by specific genetic variants. To identify potential risk genes associated with psychiatric disorders, we further obtained the risk gene sets of ASD, BD and SCZ prioritized by H-MAGMA^40^. We then estimated the overlap between the differentially expressed genes and each of these prioritized risk gene sets and identified a subset of differentially expressed genes potentially regulated by *cis*-eQTLs in frontal cortex or implicated in psychiatric disorders through chromatin interaction-informed mapping.

### Statistical analyses

All analyses and statistical tests in this study were performed using Python (version 3.9.15) and R (version 4.2.1), as described in the Methods and figure legends. The relationships among tissue-specific and region-specific pseudogenes, parent protein-coding genes, and protein-coding genes were visualized using the circlize package^71^. The distribution of different TD genes across brain regions and the annotation of functionally characterized pseudogenes were displayed using the ComplexHeatmap package^72^. Additional visualizations were made with the ggplot2 package^73^. All box plots illustrate the first and third quartiles as the lower and upper bounds, with a central band representing the median value and whiskers extending to 1.5 times the interquartile range (IQR).

## Data availability

The publicly available datasets and corresponding metadata used in this study were downloaded from Synapse (https://www.synapse.org), as listed in the **Supplementary Tables S1-S4**. The expression data of the pseudogenes and protein-coding genes in human and macaque, quantified based on the computational framework in this study, have been deposited to Zenodo (<u>10.5281/zenodo.16810204</u>). Specifically, the deposited datasets include: FPKM-normalized expression matrices of pseudogenes and protein-coding genes in non-brain tissues from human (**Supplementary Tables S6–S7**), FPKM-normalized expression matrices in diverse brain regions from both human and macaque (**Supplementary Tables S8–S11**), and raw count matrices of pseudogenes and protein-coding genes in human brain samples diagnosed with psychiatric disorders (**Supplementary Tables S14–S15**). Each matrix contains genes as rows and tissue or brain region samples as columns.

## Code availability

The custom scripts for this study were available at https://github.com/gersteinlab/brainPgene.

## Author contributions

M.G. conceived the project; Y.J. and Y.T.Y. analyzed the data; C.S., T.H. and H.W. contributed to data analysis; Y.J. and Y.T.Y. interpreted the results; Y.J. and Y.T.Y. prepared the manuscript. All authors read and approved the final manuscript.

