## Supplementary Information for "Comprehensive analysis of pseudogene expression in human and macaque brains compared with other tissues"

**Affiliation**

**Supplementary Information**

Supplementary Figures S1-S15

Supplementary Tables S1-S18

### Supplementary Figure

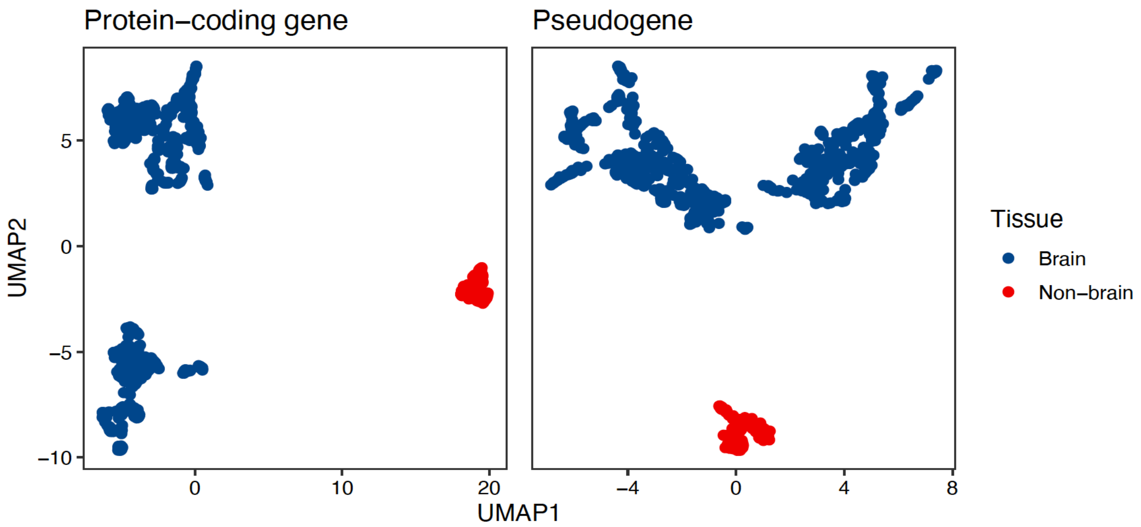


**Figure S1.** Uniform Manifold Approximation and Projection (UMAP) clustering of brain and non-brain tissues based on the gene-level expression of protein-coding genes (*left*) and pseudogenes (*right*).

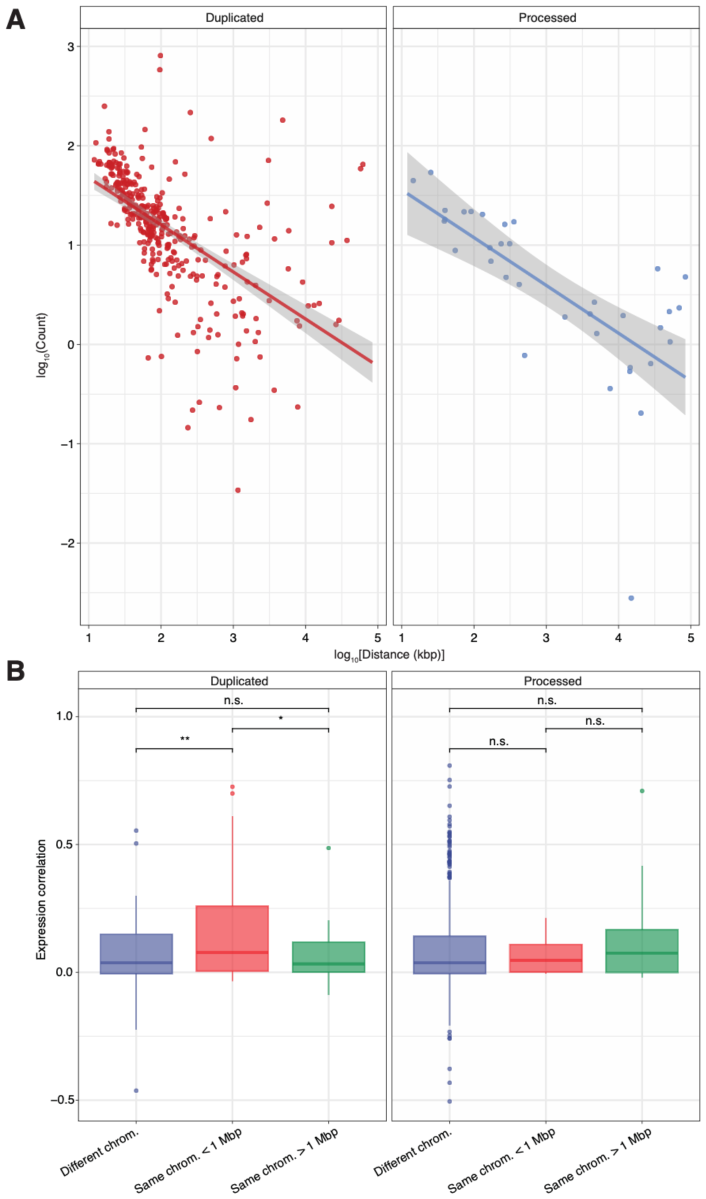


**Figure S2.** Hi-C interaction and expression correlation between pseudogenes and their parent genes. **(A)** Dot plots illustrate the inverse linear relationship between genomic distance (*x*-axis) and Hi-C count frequency (*y*-axis) for duplicated and processed pseudogenes relative to their parent genes. **(B)** Box plots display the distribution of expression correlations between duplicated and processed pseudogenes, stratified by the genomic context of the pseudogene–parent gene pair: interchromosomal (different chromosomes), intrachromosomal within 1 Mbp, and intrachromosomal greater than 1 Mbp. *, *p*-value < 0.05; **, *p*-value < 0.01; n.s., non-significant.

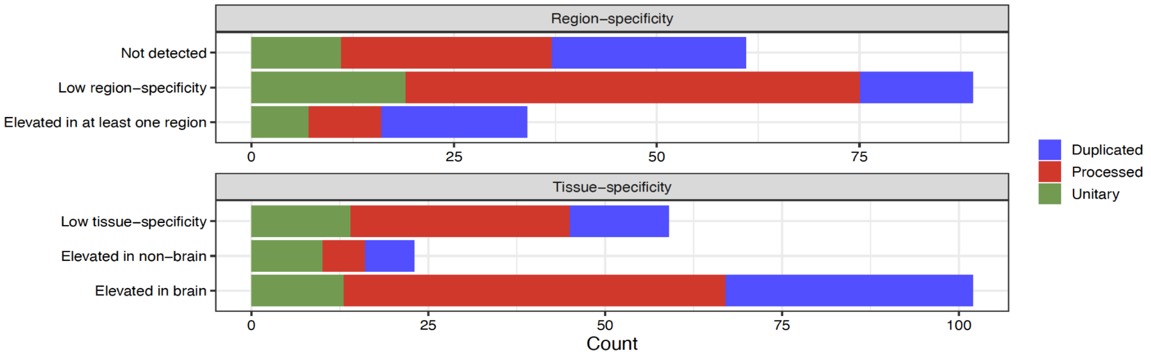


**Figure S3.** Bar plots depict the number of pseudogenes of various types identified as differentially expressed across distinct brain regions (*top*) and between brain and non-brain tissues (*bottom*).

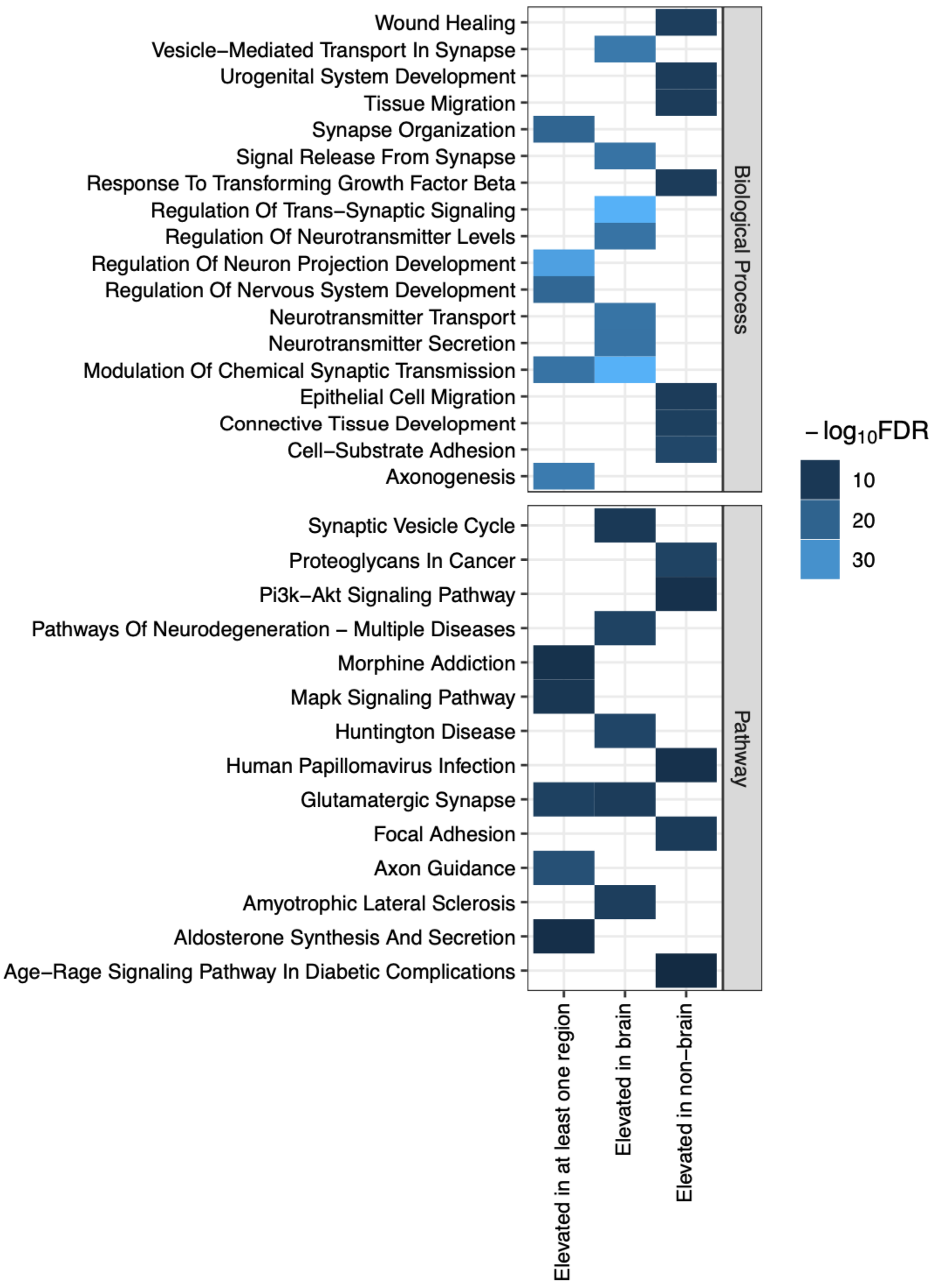


**Figure S4.** Heatmaps show the biological process (*top*) and KEGG pathway (*bottom*) enrichment analyses of protein-coding genes identified as differentially expressed across distinct brain regions and between brain and non-brain tissues.

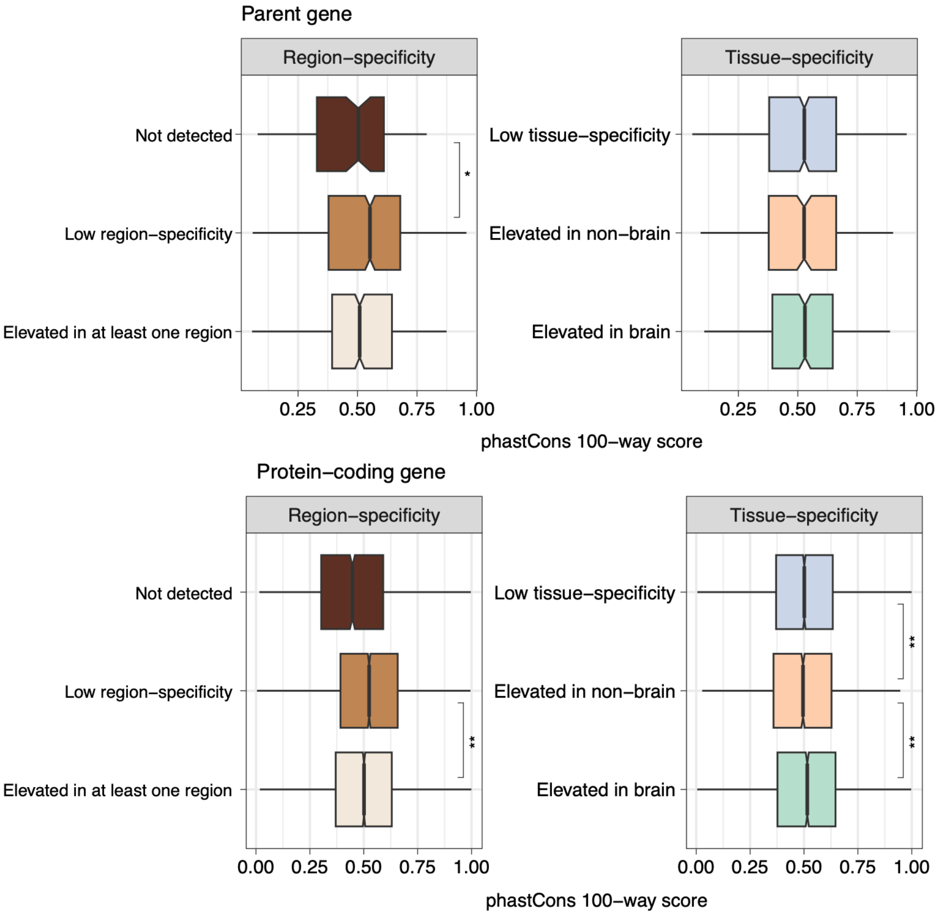


**Figure S5**. Box plots represent phastCons 100-way conservation score for region- and tissue-specific parent genes (*top*) and all protein-coding genes (*bottom*). *, *p*-value < 0.05; **, *p*-value < 0.01. Non-significant comparisons are not labeled.

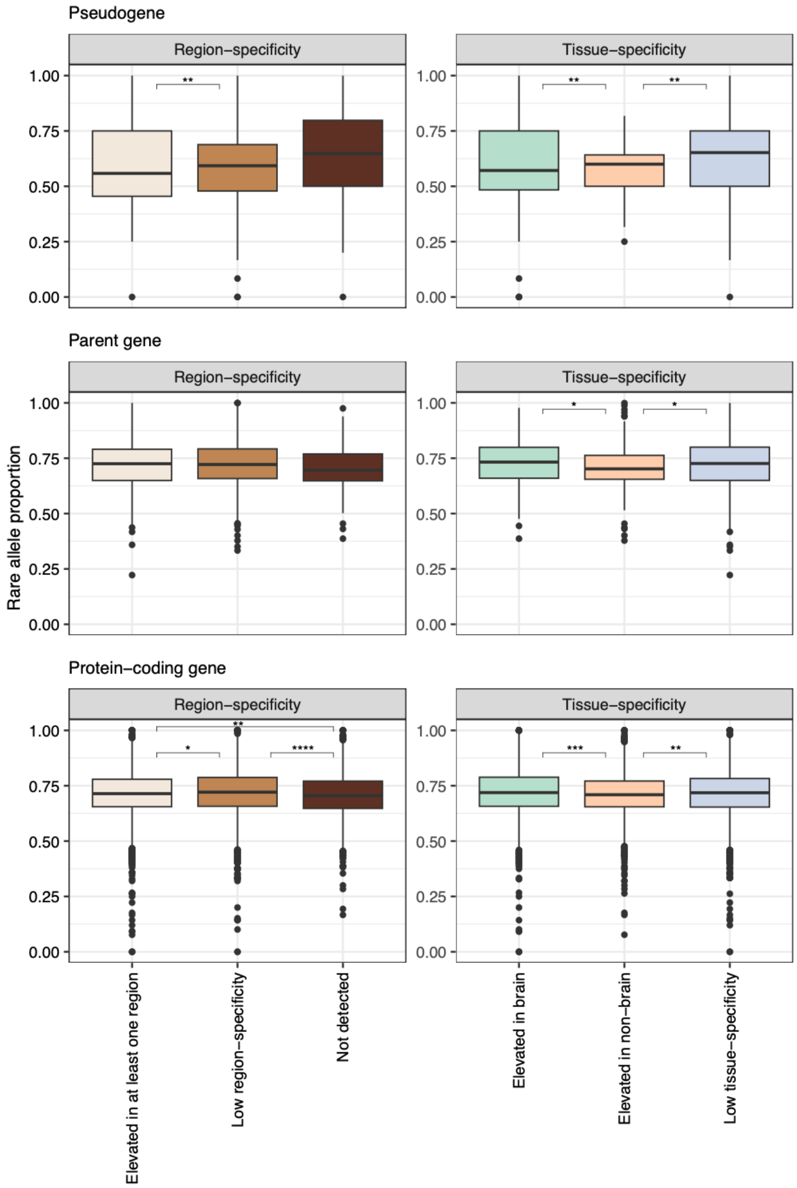


**Figure S6.** Box plots show the distribution of rare allele proportions across pseudogenes, parent genes, and protein-coding genes. Within each gene biotype, genes are further categorized based on their identification as region- or tissue-specific differentially expressed genes. *, $p$-value < 0.05; **, $p$-value < 0.01; ***, $p$-value < 0.001; ****, $p$-value < 0.0001. Non-significant comparisons are not labeled.

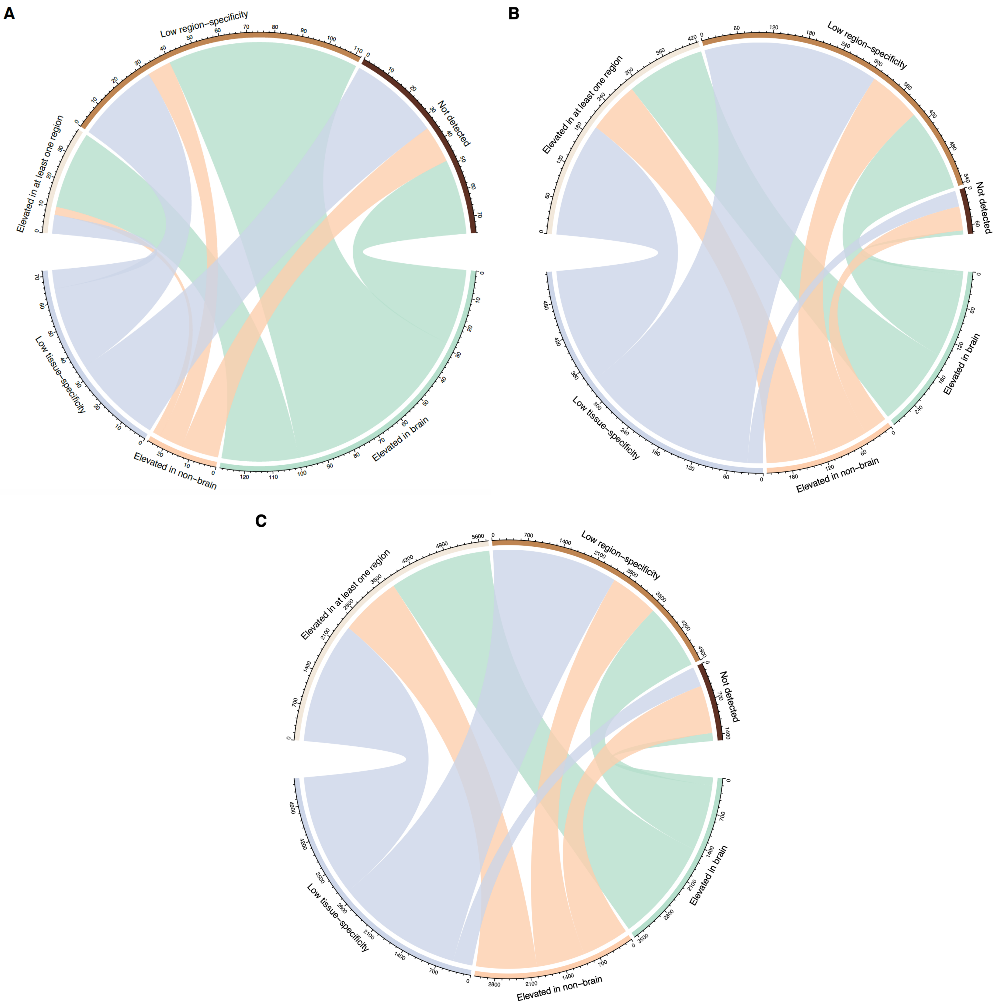


**Figure S7.** Chord diagrams show the mapping of differentially expressed pseudogenes **(A)**, parent genes **(B)**, and protein-coding genes **(C)** identified between brain and non-brain tissues to their region-specific differential expression profiles.

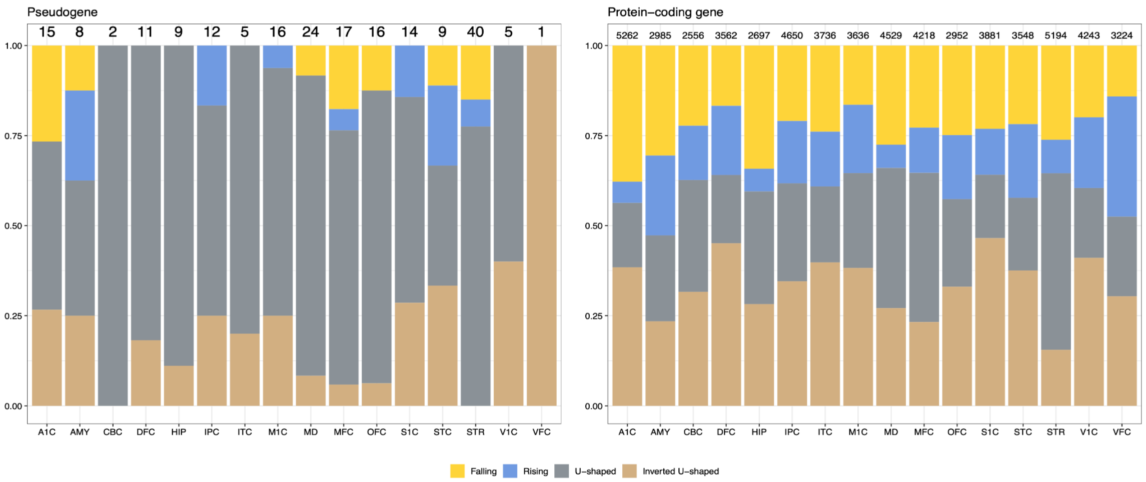


**Figure S8.** Bar plot shows the proportion of different classes of temporally dynamic pseudogenes (*left*) and protein-coding genes (*right*) across multiple macaque brain regions.

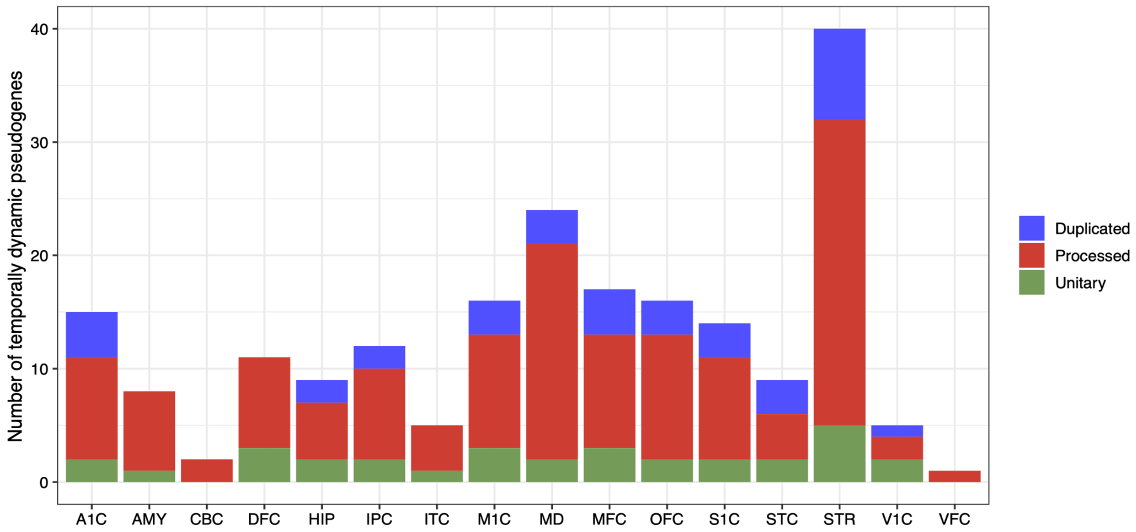


**Figure S9.** Bar plot depicts the number of temporally dynamic pseudogenes, categorized by type, across multiple macaque brain regions.

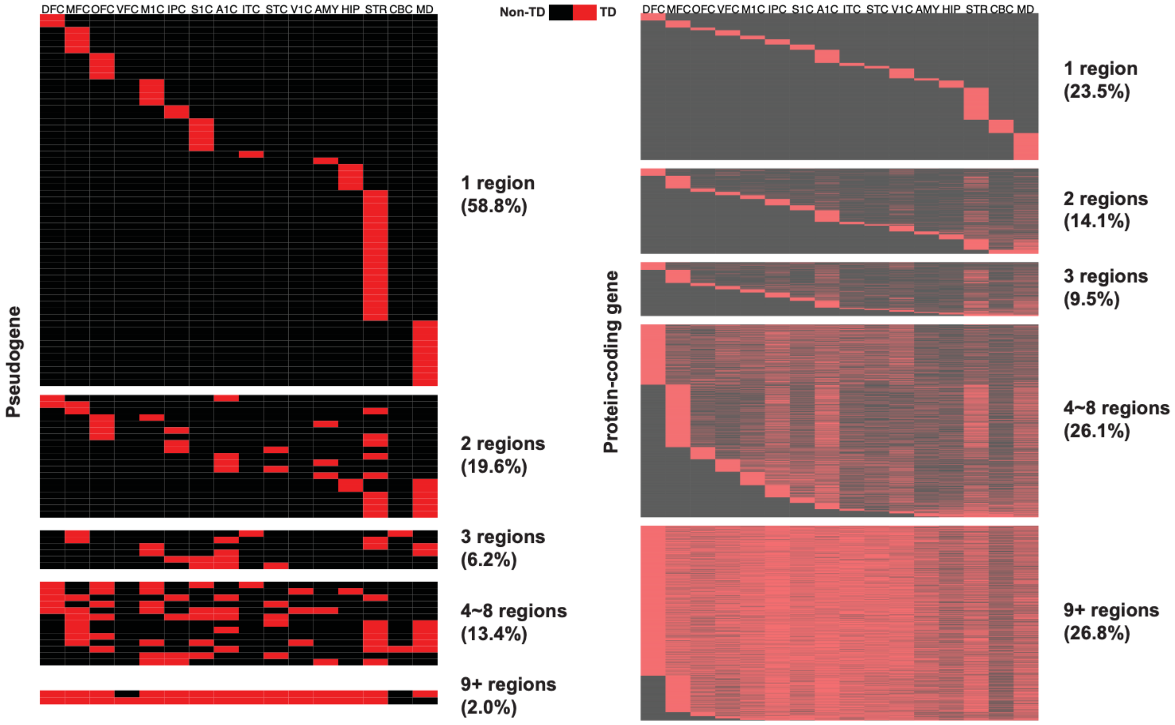


**Figure S10.** Heatmaps present temporally dynamic pseudogenes (*left*) and protein-coding genes (*right*) that are uniquely identified in a single macaque brain region or shared across multiple macaque brain regions. TD, temporally dynamic.

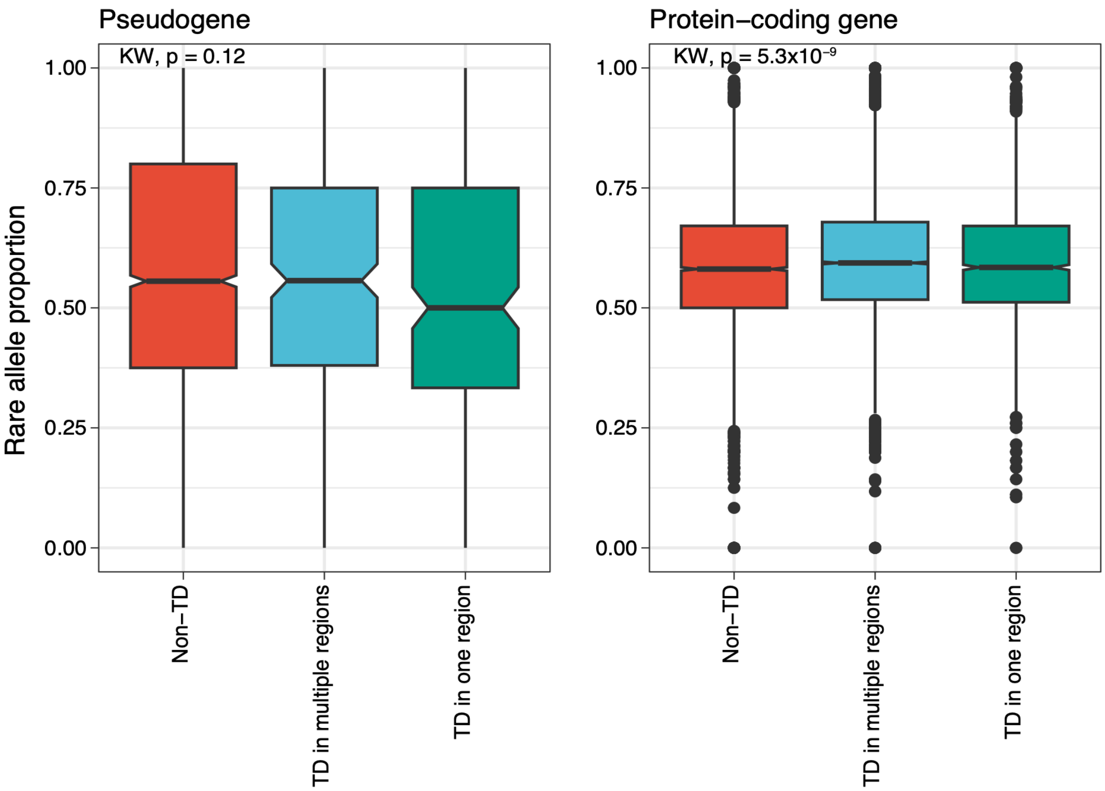


**Figure S11.** Box plots show the distribution of rare allele proportions for temporally dynamic pseudogenes (*left*) and protein-coding genes (*right*). KW, Kruskal-Wallis; TD, temporally dynamic.

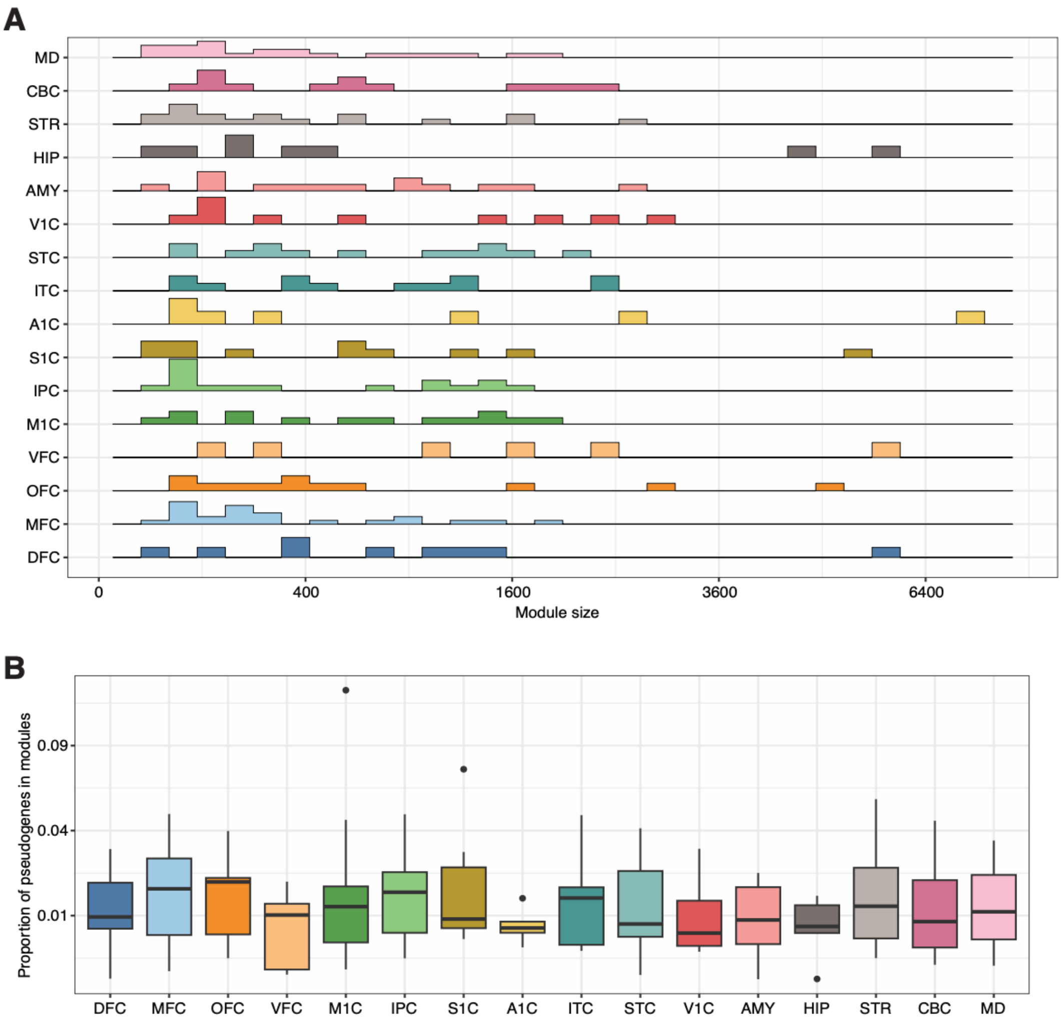


**Figure S12.** Functional inference of pseudogenes using WGCNA in the macaque brain. **(A)** Ridge plot shows the distribution of co-expression module sizes (excluding Module 0) across different brain regions. **(B)** Box plot illustrates the proportion of pseudogenes within co-expression modules (excluding Module 0) for each brain region.

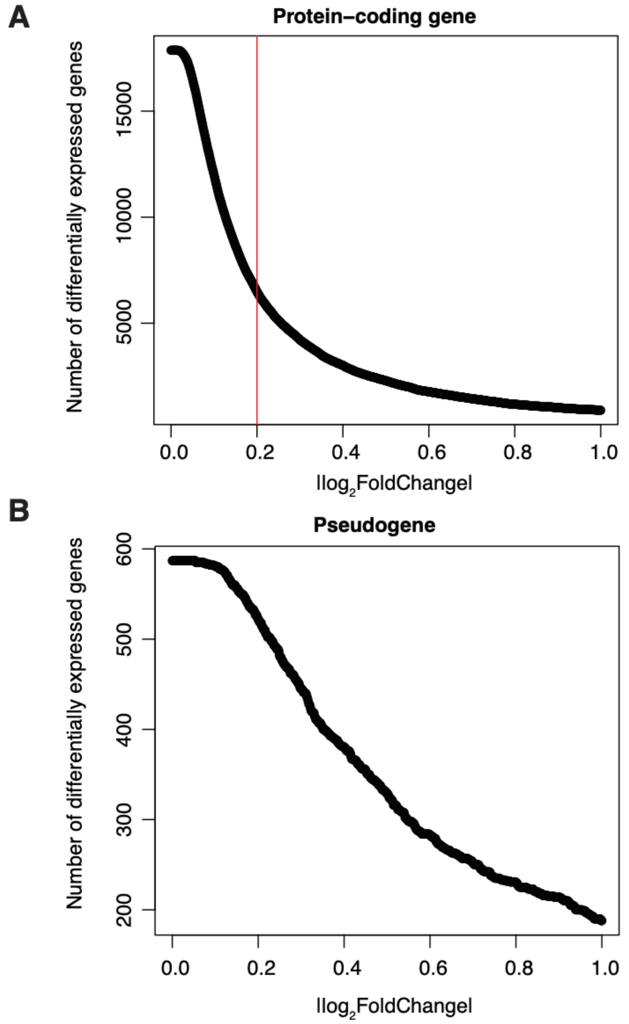


**Figure S13.** Scatter plots show the relationship between the number of differentially expressed genes and $|\log_2\text{FoldChange}|$ for protein-coding genes **(A)** and pseudogenes **(B)**. The vertical line in **(A)** indicates the threshold for protein-coding genes.

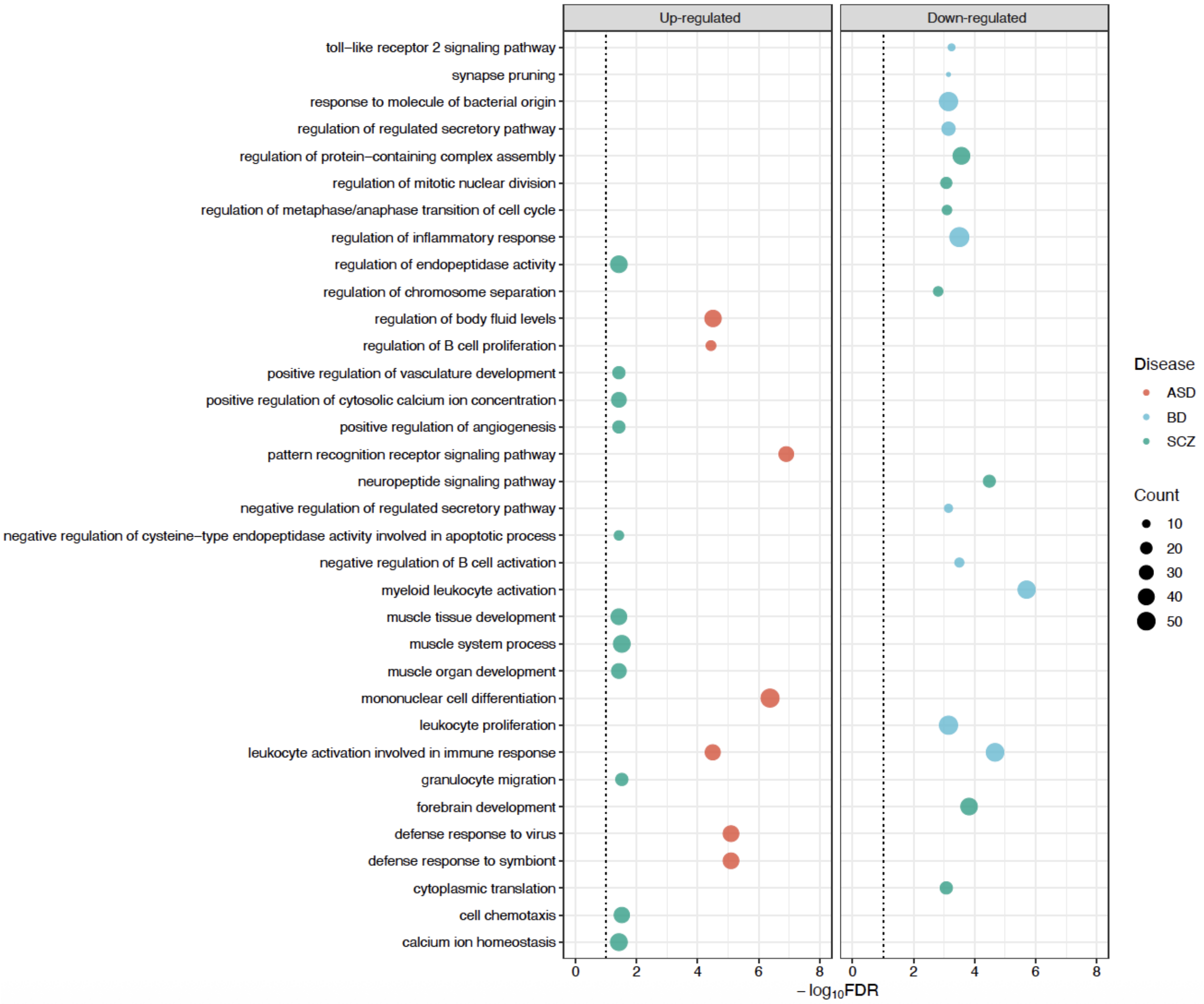


**Figure S14.** Bubble plots show enriched biological processes among upregulated and downregulated differentially expressed protein-coding genes in ASD, BD, and SCZ. The *x*-axis represents the enrichment significance as -$\log_{10}$FDR, where higher values indicate stronger enrichment. The *y*-axis lists the most significantly enriched terms. Bubbles are color-coded by disease, and their size indicates the number of differentially expressed genes associated with each term.

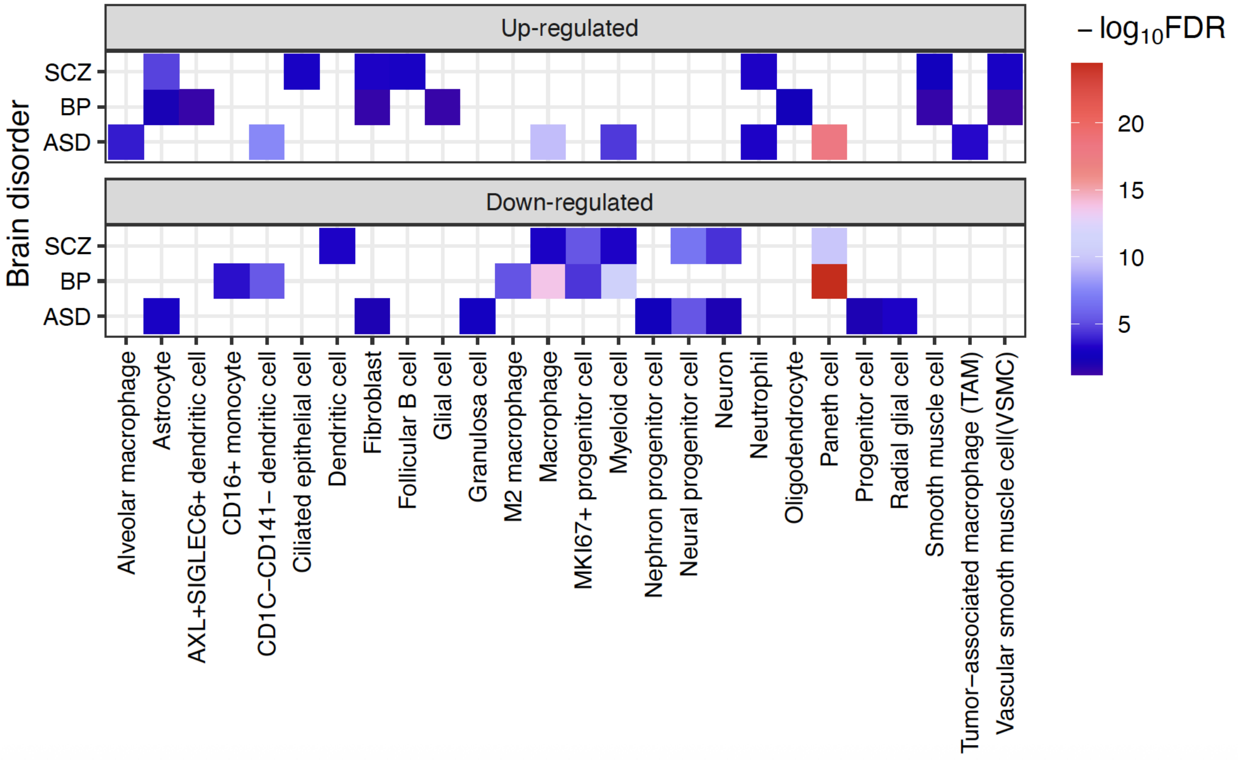


**Figure S15.** Heatmaps show the enriched cell types of upregulated and downregulated differentially expressed protein-coding genes in ASD, BD, and SCZ. The most significantly enriched cell types are shown on the *x*-axis, and brain disorders are listed on the *y*-axis. Color intensity represents the enrichment significance as $-\log_{10}$FDR.

### Supplementary Table

**Table S1.** Metadata of the brain samples from diverse brain regions that were used in this study.

**Table S2.** Metadata of the brain samples diagnosed with psychiatric disorders that were used in this study.

**Table S3.** Metadata of the macaque brain samples that were used in this study.

**Table S4.** Metadata of the human non-brain samples that were used in this study.

**Table S5.** Pseudogene annotations for the macaque genome (rheMac10) generated in this study.

**Table S6.** Expression estimates (FPKM) of pseudogenes in non-brain samples from human.

**Table S7.** Expression estimates (FPKM) of protein-coding genes in non-brain samples from human.

**Table S8.** Expression estimates (FPKM) of pseudogenes in diverse brain regions from human.

**Table S9.** Expression estimates (FPKM) of protein-coding genes in diverse brain regions from human.

**Table S10.** Expression estimates (FPKM) of pseudogenes in diverse brain regions from macaque.

**Table S11.** Expression estimates (FPKM) of protein-coding genes in diverse brain regions from macaque.

**Table S12**. Temporally dynamic genes in diverse brain regions from human.

**Table S13.** Temporally dynamic genes in diverse brain regions from macaque.

**Table S14.** Expression estimates (raw count) of pseudogenes in human brain samples diagnosed with psychiatric disorders.

**Table S15.** Expression estimates (raw count) of protein-coding genes in human brain samples diagnosed with psychiatric disorders.

**Table S16.** Differentially expressed genes identified for ASD, BD and SCZ.

**Table S17.** The disorder risk genes prioritized by H-MAGMA for ASD, BD and SCZ.

**Table S18.** The pairs between pseudogenes and their parent protein-coding genes in human genome.
